# A “suicide” BCG strain provides enhanced immunogenicity and robust protection against *Mycobacterium tuberculosis* in macaques

**DOI:** 10.1101/2023.11.22.568105

**Authors:** Alexander A Smith, Hongwei Su, Joshua Wallach, Yao Liu, Pauline Maiello, H Jacob Borish, Caylin Winchell, Andrew W Simonson, Philana Ling Lin, Mark Rodgers, Daniel Fillmore, Jennifer Sakal, Kan Lin, Valerie Vinette, Dirk Schnappinger, Sabine Ehrt, JoAnne L. Flynn

## Abstract

Intravenous (IV) BCG delivery provides robust protection against *Mycobacterium tuberculosis* (Mtb) in macaques but poses safety challenges. Here, we constructed two BCG strains (BCG-TetON-DL and BCG-TetOFF-DL) in which tetracyclines regulate two phage lysin operons. Once the lysins are expressed, these strains are killed in immunocompetent and immunocompromised mice yet induced similar immune responses and provided similar protection against Mtb challenge as wild type BCG. Lysin induction resulted in release of intracellular BCG antigens and enhanced cytokine production by macrophages. In macaques, cessation of doxycycline administration resulted in rapid elimination of BCG-TetOFF-DL. However, IV BCG-TetOFF-DL induced increased pulmonary CD4 T cell responses compared to WT BCG and provided robust protection against Mtb challenge, with sterilizing immunity in 6 of 8 macaques, compared to 2 of 8 macaques immunized with WT BCG. Thus, a “suicide” BCG strain provides an additional measure of safety when delivered intravenously and robust protection against Mtb infection.

## Introduction

*Mycobacterium bovis* BCG, also known as Bacille Calmette-Guérin or BCG, is a live *Mycobacterium bovis* strain that was attenuated by serial passaging in vitro and remains the only tuberculosis (TB) vaccine approved for use in humans. BCG protects children against miliary TB and TB meningitis but is only partially protective against pulmonary TB ^1^. Typically, BCG is delivered by intradermal injection, but studies in non-human primates revealed that endobronchial instillation or high-dose intravenous administration result in improved protection against *M. tuberculosis* (Mtb) infection ^2,3^. Mucosal and intravenous delivery of BCG pose the risk of developing disseminated BCGosis, with potentially fatal outcomes in immunocompromised individuals ^4^, such as HIV infected children ^5^, children with Mendelian susceptibility to mycobacterial disease (MSMD) ^6^ and patients with chronic granulomatous disease ^7^. Furthermore, intravesical BCG therapy is the most successful immunotherapy for bladder cancer but is also associated with severe and possibly fatal complications in approximately 5% - 8% of patients ^8–11^.

We sought to construct a BCG strain that can be killed by addition or removal of a small molecule, such as a tetracycline ^12–14^ and would be safer than BCG so that it could be used with non-conventional delivery approaches and increased dosage. Mycobacteriophages use lytic enzymes to kill their host cells including holin proteins that permeabilize the cytoplasmic membrane, endolysins that cleave peptidoglycan and lipid hydrolases/esterases that degrade the outer membrane ^15^. We took advantage of the lysin operons from the mycobacteriophages D29 and L5 ^16^ that encode a holin, an endolysin (lysin A) and a mycolylarabinogalactan esterase (lysin B) to construct regulated kill switches for BCG. We hypothesized that killing by inducing cell lysis may make BCG more immunogenic due to the release of intracellular antigens. In a previous study 5 x 10^7^ CFU of BCG (SSI strain) were delivered intravenously to rhesus macaques which resulted in substantial (∼10,000 fold) reduction of live Mtb in the animals by 4 weeks, with 6 of 10 macaques showing sterile protection ^2^. Here, we assessed whether doxycycline-controlled lysins can kill BCG (Pasteur strain) in mice and macaques. We evaluated the level of persistence of BCG kill switch strains and the induced immune responses, and whether these strains administered intravenously could provide protection against Mtb challenge in mice and macaques.

## Results

### Construction and in vitro characterization of BCG kill switch strains

We evaluated the lysin operons of the mycobacteriophages L5 (L5L) and D29 (D29L) to construct BCG strains that could be efficiently killed by adding or removing tetracyclines such as anhydrotetracycline (atc) or doxycycline (doxy). We first tested functionality of these lysin operons in BCG using a TetON expression system ^12^. Lysin transcription is repressed by a wild-type (WT) tetracycline repressor (TetR) that binds to a WT tetO positioned between the −10 and −35 elements of the lysin promoter. Addition of atc/doxy inactivates TetR and turns on lysin expression (**Fig. S1a**). As expected, atc addition prevented growth of BCG strains that carried TetON-D29L or TetON-L5L integrated into the chromosome, although escape mutants prevented complete killing and resulted in regrowth in liquid culture (**Fig1a, Fig. S1b**). The optical densities (OD) of BCG cultures carrying both TetON-D29L and TetON-L5L (BCG TetON-DL) rapidly declined when atc was added indicating that death occurred by cell lysis (**Fig. 1b**, **Fig. S2a**). The impact of atc on OD was similar regardless at what time after inoculation it was added. Cytosolic enolase and proteasome subunit B accumulated in the culture filtrate of BCG TetON-DL following treatment with atc, which demonstrated that atc-induced lysin expression indeed resulted in bacterial lysis (**Fig. S2b**). The combined induction of two lysins reduced the fraction of escape mutants by almost two orders of magnitude compared to those observed with single lysin strains (**Fig. S2c**).

**Figure 1.**
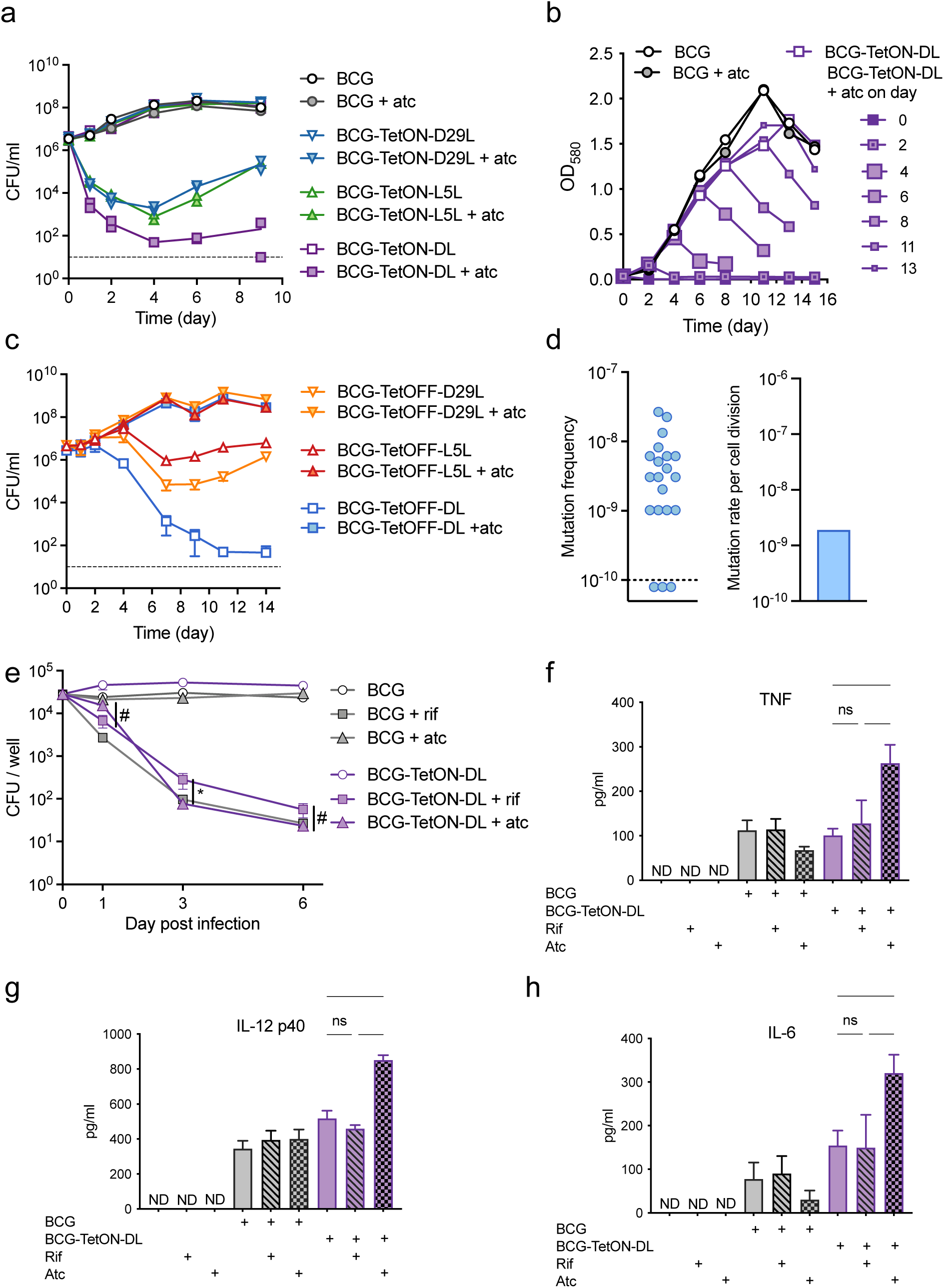
Induction of phage lysins results in cell lysis with low frequency of escape and promotes macrophage proinflammatory cytokine production. (a) Growth of BCG TetON single and dual lysin kill switch strains with and without atc. Individual CFU data points from two replicate cultures are depicted. (b) Impact of BCG TetON dual lysin kill switch induction at different times of growth. (c) Growth of BCG TetOFF single and dual lysin kill switch strains with and without atc. Data are means ± SD from triplicate cultures. Error bars are frequently too small to be seen. (d) Fraction of resistant mutants per culture and mutation rate per cell division of BCG-TetOFF-DL. The resistance rate in 20 individual cultures was determined in a fluctuation assay. The number of mutations per cell division was calculated using the bz-rates web-tool. (e) CFU quantification of WT BCG and BCG-TetON-DL (plus or minus atc or rifampin) during infection of murine BMDMs. Data are means ± SD from triplicate cultures. Multiple unpaired t-tests were performed on log_10_-transformed data comparing BCG-TetON-DL treated with rif to BCG-TetON-DL treated with atc at each time point with Holm-Šídák adjusted p-values. * p < 0.05, # 0.05 < p < 0.10. (f, g, h) Quantification of TNF (f), IL-12 p40 (g), and IL-6 (h) production by macrophages infected with BCG or BCG-TetON-DL and treated with rifampin or atc for 20 hrs. Data are means ± SD from triplicate cultures. Significance was determined by one-way ANOVA with Tukey’s adjusted p-values. **** p < 0.0001, ***, p < 0.001, ** p < 0.01, * p < 0.05, ns p > 0.1. ND, not detected. All data are representative of two or three independent experiments.

To test whether L5L/D29L could also be used to generate atc/doxy-addicted BCG, which would require a tetracycline to grow, we cloned the lysin operons into two expression systems that are both silenced by atc/doxy. L5L was cloned into a TetPipOFF system ^17^, in which WT TetR represses a second transcriptional repressor, PipR, that in turn represses L5L. Here, atc inactivates TetR, which turns lysin expression off via production of PipR. D29L was cloned into an expression system controlled by a reverse TetR (revTetR), which requires atc/doxy to efficiently bind to its operator DNA (tetO_4C5G_) ^13^. Heterodimerization of TetR and revTetR produced in the same cell was prevented by using single-chain versions of each of these repressors ^18^. Both BCG-TetOFF lysin strains were killed when atc was removed from the cultures (**Fig. 1c**, **Fig S2d**). The onset of killing was delayed compared to the TetON strains, which is likely due to the time required to dilute intracellular atc from the bacteria. Death of the BCG-TetOFF strains was accompanied by cell lysis (**Fig. S2b**), and expression of two lysins led to enhanced killing and reduced the fraction of escape mutants (**Fig. S2e**). Analysis of the dual TetOFF lysin strain, BCG-TetOFF-DL, with fluctuation assays detected ∼2 x 10^-9^ escape mutants per cell division (**Fig. 1d**).

### Lysin induction in intracellular BCG promotes proinflammatory cytokine production

We infected bone marrow derived mouse macrophages (BMDM) with BCG and BCG-TetON-DL and treated them with rifampin (RIF) or atc (**Fig. 1e**). RIF killed both intracellular BCG and BCG-TetON-DL effectively. Atc had no effect on BCG but killed BCG-TetON-DL presumably via lysin induction. BCG and BCG-TetON-DL both induced TNF, IL12 p40 and IL-6 production by BMDMs. However, cytokine production was significantly increased when killing was mediated by phage lysin expression (**Fig. 1f-h**). These data suggest that intracellular lysis of BCG-TetON-DL enhanced cytokine production compared to that stimulated by live BCG or by BCG and BCG- TetON-DL that had been killed by RIF.

### Lysin induction kills BCG in immune competent and immune deficient mice

In C57BL/6 mice BCG-TetON-DL established infection in lungs and spleens and persisted similar to BCG following intravenous injection **(Fig. 2 a,b**). When mice were treated with doxy starting seven days post infection, BCG-TetON-DL was killed in lungs and spleen, while doxy had a modest impact on WT BCG. BCG-TetOFF-DL similarly established infection in C57BL/6 mice that received doxy containing chow (**Fig. 2c,d**); its titers declined slowly in the lungs even when mice received doxy, likely due to doxy levels that were insufficient to maintain persistence in the context of host immunity, but remained stable in spleens. When doxy was eliminated from the mouse chow starting 14 days post infection, BCG-TetOFF-DL lost viability in lungs and spleens and was cleared from the lungs within 6 weeks of doxy withdrawal.

**Figure 2.**
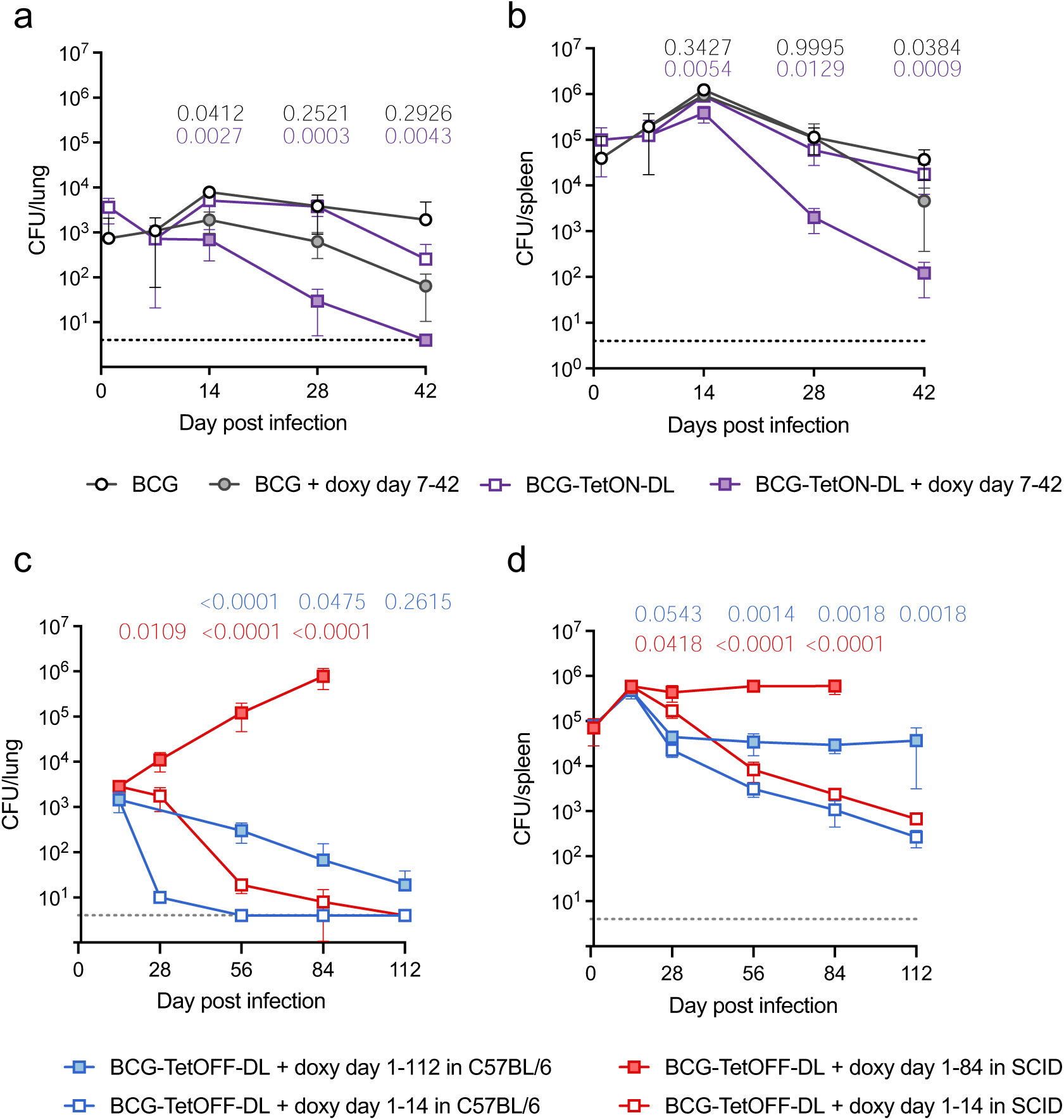
Phage lysin induction kills BCG in immune competent and immune deficient mice. (a, b) CFU quantification from lungs (a) and spleens (b) of BCG and BCG-TetON-DL infected C57BL/6 mice treated or not with doxycycline (doxy) starting day 7 post infection. Data are means ± SD from 3-5 mice per group and time point. (c, d) CFU quantification from lungs (c) and spleens (d) of BCG-TetOFF-DL infected C57BL/6 and SCID mice treated or not with doxy for the indicated times. Data are means ± SD from 4-5 mice per group and time point. Multiple unpaired t-tests run on log_10_-transformed data at each time point with Holm-Šídák adjusted p- values shown comparing BCG with BCG + doxy (black p-values), BCG-TetON-DL with BCG-TetON-DL + doxy (purple p-values), BCG-TetOFF-DL with BCG-TetOFF-DL + doxy in C57BL/6 (blue p-values) and in SCID (red p-values) mice.

To assess safety of BCG-TetOFF-DL, we infected immunocompromised SCID mice (**Fig. 2c,d**). In SCID mice that received doxy, BCG-TetOFF-DL replicated in lungs during the 84 day-long infection. In spleens the strain replicated until it reached a titer of ∼10^6^ and then persisted at a 10-fold higher burden than in C57BL/6 mice. In SCID mice that received doxy only for 14 days, BCG-TetOFF-DL was cleared from the lungs although slower than in immunocompetent C57BL/6 mice (**Fig. 2c**). In spleens, BCG-TetOFF-DL titers declined with kinetics like those observed in C57BL/6 mice (**Fig. 2d**). In a repeat experiment, BCG-TetOFF-DL was similarly eliminated from lungs and declined in viability in spleens in SCID mice that received doxy for only 14 days (Fig. S3c). Although the strain replicated less in lungs of SCID mice that received doxy for 84 weeks than in the first experiment, the data reproducibly show that BCG-TetOFF-DL does not replicate in SCID mice in the absence of doxy. We assessed lung pathology and found rare, small cellular aggregates in both groups of mice without overt signs of disease (**Fig. S3d,e**). These data demonstrate that the dual lysin BCG kill switch strains recapitulate vaccination with BCG but can be killed by lysin expression via doxy administration or withdrawal. We have no evidence for the emergence of escape mutants in any of the animal experiments and BCG-TetOFF-DL proved to be safe in immunocompromised SCID mice.

### BCG kill switch strains provide similar protection as wild type BCG against Mtb infection in mice

We vaccinated C57BL/6 mice by intravenous administration of 1x10^6^ BCG and BCG-TetOFF-DL and measured CFU in lungs and spleens 21, 56 and 84 days post vaccination (**Fig. S3a**). The mice did not receive any doxy, so that lysin expression was induced in BCG-TetOFF-DL soon after infection. Both strains lost viability in lungs, but BCG-TetOFF-DL was eliminated faster than BCG and cleared from the lungs by 56 days post vaccination. In spleens, BCG persisted at approximately 2x10^4^ CFU, while BCG-TetOFF-DL steadily lost viability, with 20 CFU remaining on day 84.

We examined pulmonary T cell responses in vaccinated mice and in mice injected with PBS (**Fig. 3a-e**). BCG and BCG-TetOFF-DL elicited effector memory CD4 T cells (**Fig. 3a**) and CD8 T cells (**Fig. 3b**) whose frequencies were reduced on days 56 and 84 in BCG-TetOFF-DL vaccinated mice compared to BCG vaccinated mice. CD153 expressed on CD4 T cells has been identified as immune mediator of host protection against Mtb infection ^19,20^. In mice vaccinated with BCG or BCG-TetOFF-DL, CD153 positive CD4 T cells were similarly enriched in the lungs on day 56 and 84 post vaccination (**Fig. 3c**). Lung resident memory CD4 T cells were also induced by vaccination, although the responses in mice vaccinated with BCG-TetOFF-DL were slightly lower than in mice vaccinated with BCG (**Fig. 3d**). Finally, we measured the frequency of pulmonary cytokine (TNF, IFNγ, IL2, IL17A) expressing CD4 T cells and observed similar responses in BCG and BCG-TetOFF-DL vaccinated mice (**Fig. 3e**). Collectively these data indicate that vaccination with BCG-TetOFF-DL resulted in robust pulmonary T cell responses that were similar to those elicited by BCG vaccination, despite the absence of doxy from the beginning of vaccination and faster clearance of BCG-TetOFF-DL from the lungs (**Fig. S3a**).

**Figure 3.**
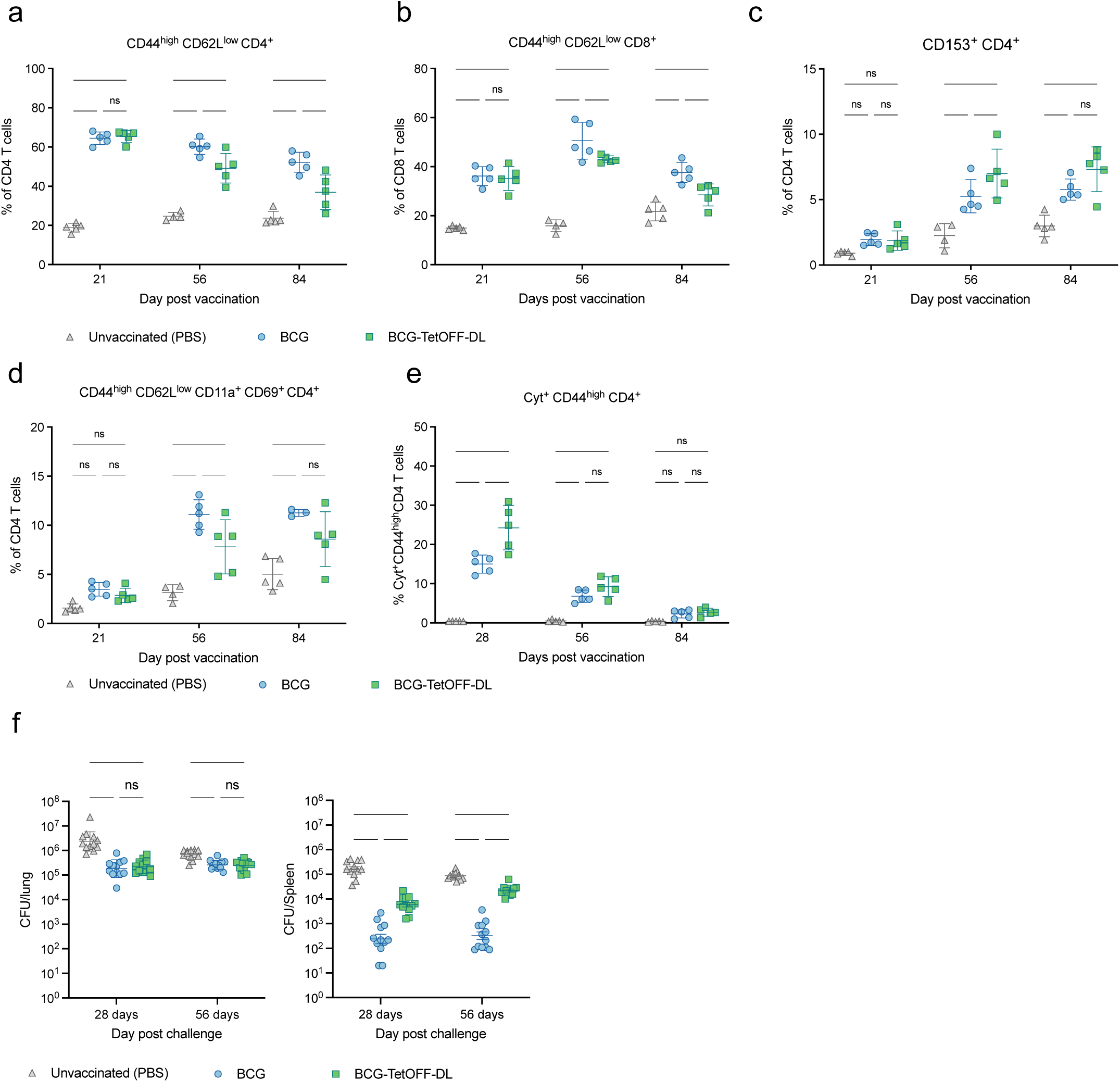
BCG-TetOFF-DL and wt BCG provide similar protection against Mtb infection in mice. **Mice were** i.v. vaccinated with BCG or BCG-TetOFF-DL or received PBS. (a-e) Quantification of T cell subsets in mouse lungs from mice vaccinated with BCG or BCG-TetOFF-DL; (a) effector memory CD4 T cells; (b) of effector memory CD8 T cells; (c) CD153 expressing CD4 T cells; (d) lung resident memory CD4 T cells; (e) Cytokine (TNF, IFNγ, IL2, IL17A) expressing lung cells following ex vivo restimulation with PPD prior to intracellular cytokine staining and Boolean OR gating. Symbols are data from individual mice with lines indicating mean ± SD. Two-way ANOVA was performed, and Tukey’s adjusted p-values are shown for each time point. **** p < 0.0001, ***, p < 0.001, ** p < 0.01, * p < 0.05, ns p > 0.10. (f) Bacterial burden in lungs and spleens of vaccinated and PBS treated mice. Mice were infected with Mtb H37Rv by aerosol 90 days post vaccination. CFU on day one post infection were 90 ± 17. Symbols represent data from individual mice with lines indicating mean ± SEM. Two-way ANOVA was performed on log_10_- transformed data and Tukey’s adjusted p-values are shown for each time point. **** p < 0.0001, ***, p < 0.001, ** p < 0.01, * p < 0.05, ns p > 0.10.

We challenged unvaccinated and vaccinated mice with Mtb H37Rv via aerosol infection (**Fig. 3f**). Mtb H37Rv carried a hygromycin resistance cassette that allowed us to specifically detect Mtb in BCG and BCG-TetOFF- DL vaccinated mice. On day 28 post aerosol challenge, Mtb H37Rv had replicated to a mean of 4 x 10^6^ CFU in the lungs and 2 x 10^5^ CFU in the spleen of unvaccinated mice. In mice vaccinated with BCG or BCG-TetOFF- DL, the Mtb titers were reduced by approximately 10-fold (2.5 x 10^5^) in lungs. In spleens BCG vaccination reduced Mtb burden by more than 100-fold compared to unvaccinated mice, while vaccination with BCG- TetOFF-DL led to a 30-fold reduction. We attribute this difference in protection to the significantly reduced persistence of BCG-Tet-OFF-DL compared to BCG in mouse spleens (**Fig. 2**). On day 56 post challenge, both BCG strains provided reduced protection against Mtb in lungs than on day 28, while levels of protection were largely maintained in the spleens. We repeated the challenge experiment with BCG-TetON-DL with similar outcomes although there was no difference in protective efficacy provided by BCG and BCG-TetON-DL in the spleens (**Fig. S4**). Together, these data demonstrate that BCG kill switch strains protect against Mtb infection in mice comparably to BCG, although they are eliminated more rapidly than BCG from lungs and spleens in the mouse model (**Fig. 2**).

### BCG persistence study design for NHPs

We used BCG-TetOFF-DL to assess persistence in non-human primates. To determine whether the doxycycline-regulated lysin expression was functional in vivo in macaques and whether BCG-TetOFF-DL persisted or was diminished after removal of doxy treatment, we administered 5x10^7^ BCG-TetOFF-DL CFU IV to nine Mauritian cynomolgus macaques (MCM)(**Fig. 4a**). Group A was given daily doxy beginning one day prior to BCG inoculation and continued for 2 weeks and was euthanized at 4 weeks post-vaccination. Group B had the same 2-week doxy regimen and was euthanized at 8 weeks. Group C had an 8-week doxy regimen and was euthanized at 8 weeks. Doxy administration was expected to prevent the expression of the two lysin genes, maintaining the ability of BCG-TetOFF-DL to persist and/or grow in the macaques. Withdrawing doxy treatment after two weeks in groups A and B was expected to result in expression of the lysin genes, preventing replication or persistence of BCG-TetOFF-DL. Group C was maintained on doxy for the full 8 weeks of the study as a control group. We performed a bronchoalveolar lavage (BAL) 4 weeks post-vaccination. At necropsy, we plated tissues for BCG-TetOFF-DL on plates containing atc. Single cell suspensions of BAL and tissue samples were assessed via flow cytometry for immune responses induced by BCG.

**Figure 4.**
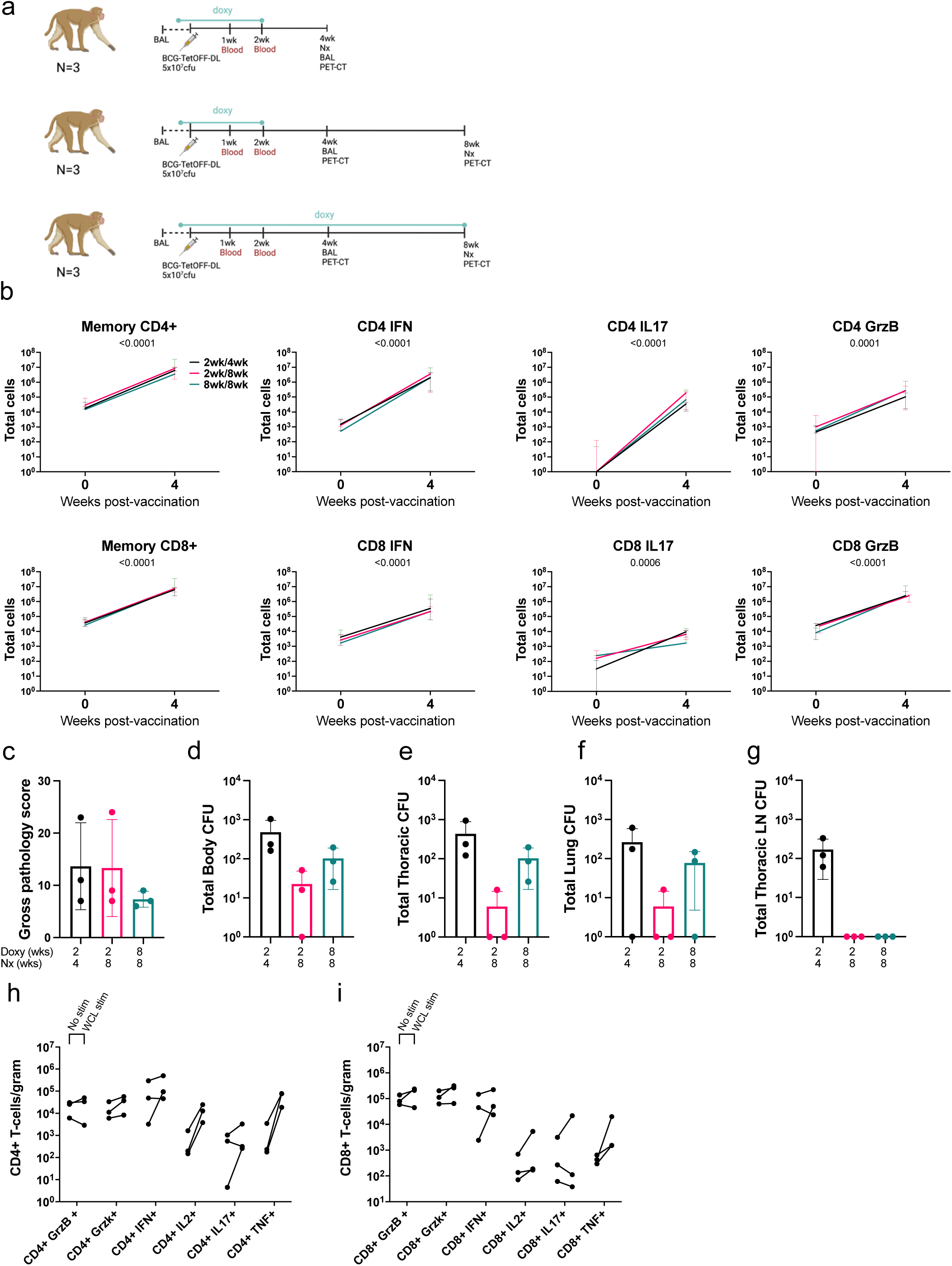
Discontinuation of doxy limits BCG-TetOFF-DL persistence in NHP. (a) Persistence study design (b) Memory CD4+ pre vaccination, memory effector CD4+ 4 weeks post vaccination, memory CD8+ pre vaccination and memory CD8+ 4 weeks post vaccination (n=3/group) total cell numbers isolated from the BAL, stimulated with H37Rv WCL.Two-way ANOVA was performed. There were no treatment effects, but total cells increased over time in all cell types. Median and range shown. (c) Gross pathology scores at necropsy. (d-g) Bacterial burden (CFU) of BCG-TetOFF-DL CFU recovered from total body of animal (d), thoracic cavity (e), lungs (f) and thoracic lymph nodes (g) (n=3/group, mean and SD shown). Number of effector CD4+ cells (h) and CD8+ cells (i) per gram of lung tissue, stimulated with media or H37Rv WCL (average of 4 lung tissues per animal, n=3 animals/group). Memory is defined as CD45RA+CD28-, CD45RA-CD28+ or CD45RA-CD28-.

### BCG-TetOFF-DL induced robust T cell responses in airways

After IV BCG-TetOFF-DL vaccination, the number of memory (CD45RA+CD28-, CD45RA-CD28+ or CD45RA- CD28-) CD4+ and CD8+ T cells recovered from the airways via BAL increased ∼100-fold in all three groups, which is indicative of the generation of an IV BCG-dependent vaccine response, based on our previous studies ^2^ (**Fig. 4b**). This corresponded with an increase in the number of cytokine and cytotoxic molecule producing effector T cells in the BAL (**Fig. 4b**). Cell numbers and effector molecule expression between groups remained consistent, suggesting the duration of the doxy regimen and time of necropsy did not play a significant role in the generation of the immune response (**Fig 4h-i**). These data suggest that a BCG-dependent, multi-faceted, immune response was generated in all animal groups within 4 weeks post-vaccination. Stimulation with *M. tuberculosis* H37Rv whole-cell lysate did not have a large influence on the number of cytokine producing CD4+/CD8+ T-cells, when compared to unstimulated controls. This is likely due to the systemic spread of BCG- TetOFF-DL when administered via the intravenous route, resulting in common mycobacterial antigens persisting at this early time point post-vaccination.

### Cessation of doxycycline reduced BCG-TetOFF-DL bacterial burden in macaque tissues

Gross pathology score is a quantitative measure of grossly apparent mycobacterial related lesions at necropsy. It takes into account granuloma numbers and lung lobes involved, lymph node and spleen size and granuloma involvement and any other evidence of infection ^21^. Gross pathology scores (**Fig. 4c**) were relatively low in all groups, which was expected since the animals were not challenged with Mtb. Two animals in group A, and 1 animal each in groups B and C had a few small granulomas found at necropsy, with 1 granuloma in a group A animal and 1 granuloma in a group B animal positive for BCG-TetOFF-DL CFU. BCG-TetOFF-DL bacterial burden was assessed in multiple tissue samples to assess the efficiency of the doxy dependent self-killing of the BCG-TetOFF-DL strain in all lung lobes, thoracic and peripheral lymph nodes and extrapulmonary organs (spleen and liver) of each animal (**Fig. 4d-g**). BCG-TetOFF-DL CFU was recovered at low levels from the lungs in 2 of 3 animals in group A (doxy stopped at 2 weeks and necropsied 4 weeks post-vaccination), from thoracic lymph nodes of all 3 animals in group A, from peripheral lymph nodes in one animal and spleen from a different animal. For group B animals (necropsied at 8 weeks post BCG-TetOFF-DL which was 6 weeks post-doxy cessation), one was sterile (no CFU recovered), and BCG-TetOFF-DL was recovered from one lung sample in one animal and from the spleen in a different animal, all at low bacterial burdens. For group C animals (treated with doxy throughout the study and necropsied at 8 weeks post-BCG-TetOFF-DL), CFU were recovered from lung lobes in two animals and from a peripheral lymph node in two animals. No culturable bacteria were recovered from the blood at weeks 1 and 2 post vaccination or in the sternum, kidneys or liver of any animals. These data indicate that killing of BCG was accelerated when doxy treatment was stopped at 2 weeks post-vaccination. Group A (doxy treatment for 2 weeks, necropsied at 4 weeks) had a mean total body CFU of 479 (range 160-1041); Group B (doxy treatment for 2 weeks, necropsied at 8 weeks) mean total body CFU was 22 (range 0-50)(**Fig. 4d**). Group C (doxy for 8 weeks, necropsy at 8 weeks) had a mean total body CFU of 102 (range 25-195)(**Fig. 4d**). Thus, there was on average a 4.5-fold reduction in CFU (p=0.2, likely influenced by small sample size) in the animals that were treated with doxy for 2 weeks vs 8 weeks and necropsied at 8 weeks (i.e. comparing groups B and C). However, it is clear that even with continuation of doxcycyline for the 8 weeks of this study (group C), BCG-TetOFF-DL bacterial burden was relatively low. This is consistent with our previous data on wild type SSI BCG ^2^. Although the animals were vaccinated with >10^7^ CFU, BCG and BCG-TetOFF-DL appear to be rapidly reduced in macaques (∼10,000 fold reduction by 8 weeks), suggesting minimal replication and/or enhanced bacterial killing.

Non-necrotizing ‘microgranulomas’ and lymphohistiocytic aggregates were seen throughout the spleen (**Fig. S5a**), thoracic lymph nodes (**Fig. S5b**) and livers (**Fig. S5c**) of animals in all groups. BCG-TetOFF-DL CFU was recovered from spleen in only 2/9 animals, liver in 0/9 animals and lymph nodes (peripheral or thoracic) in 6/9 animals.

### Lymphocyte proportions in lung and lymph nodes were similar from each group post BCG-TetOFF-DL vaccination

We compared lymphocyte composition in the lung and thoracic lymph nodes in tissue samples collected at necropsy (**Fig. S5d,e**). Expected levels of animal-to-animal variation was seen in cellular composition, however, populations remained consistent within and across groups. CD4^-^CD8^-^ (double negative) T cells were more prominent in group A lung tissues compared to group B and C, however other populations were similar. Few B cells (CD20+) were found in lung tissue compared to lymph nodes, but a higher proportion of γδ T-cells and NK cells were present in lung tissue. Cellular populations were similar across all animal groups in thoracic lymph nodes, with CD4+ T-cells being the most prominent cell type.

### T-cell responses in the lung and lymph nodes at necropsy

Comparing CD4+ and CD8+ T cell numbers in lung tissue at necropsy shows comparable levels of cells in all three groups (**Fig. 4h,i** – No stim). Similar cell numbers from animals on a short doxy regimen and necropsied at 4 weeks (Group A), and from those necropsied at 8 weeks (Group C) on a longer doxy regimen suggest a robust, multi-cellular immune response is generated and resides in the lungs within 4 weeks post vaccination with IV- BCG-TetOFF-DL. Cytokine and cytotoxic molecule producing cells were similar between all 3 groups, further reinforcing the multi-faceted response in lung tissue (**Fig. 4h,i**). A high number of CD4+ cells responded to H37Rv whole cell lysate (WCL) stimulation, resulting in the production of TNF, IFN-γ and IL-2 in the lung tissue. Cytotoxic molecules (granzyme B and K) are preformed molecules, therefore stimulation with H37Rv WCL has minimal effect on the quantities detected compared to unstimulated cells. High levels of these cytotoxic molecules were detected in all groups, in both unstimulated and stimulated samples.

In summary, IV BCG-TetOFF-DL vaccination was able to generate a robust, multi-faceted immune response comprising of effector CD4+ and CD8+ T-cells within 4 weeks post vaccination. The ‘kill-switch’ was successfully induced in a NHP model by removal of doxy, with a trend of reduction in BCG CFU 6 weeks after cessation of doxy administration. Gross pathology scores at necropsy remained low, suggesting minimal negative side-effects of IV BCG-TetOFF-DL vaccination.

### Determining protective efficacy of BCG-TetOFF-DL in macaques against Mtb infection and disease

Following the persistence study, we performed an Mtb challenge study to assess the protective efficacy of BCG- TetOFF-DL and WT BCG Pasteur in NHPs. Protection was assessed by monitoring disease progression over time using PET-CT and quantifying Mtb bacterial burden at necropsy. The immune response generated to each vaccine strain – BCG-TetOFF-DL and WT BCG Pasteur- was characterized over the duration of the study and at necropsy.

We administered 5x10^7^ BCG-TetOFF-DL or WT BCG Pasteur CFU to 8 MCMs per group, both intravenously (**Fig. 5a**). Two MCMs were unvaccinated as concurrent controls for this study; 8 additional historical unvaccinated MCM controls with the same time point for necropsy were included in this study, resulting in 10 unvaccinated control MCMs total. To reduce the possibility of confounding effects, both vaccine groups were given a 2 week doxy regimen post-vaccination which should not affect WT BCG. At 22 weeks post vaccination, we challenged all groups with 9-16 CFU Mtb strain Erdman via intrabronchial instillation (**Fig. 5a**), monitored disease progression over time using PET/CT, and necropsied 12 weeks post-challenge.

**Figure 5.**
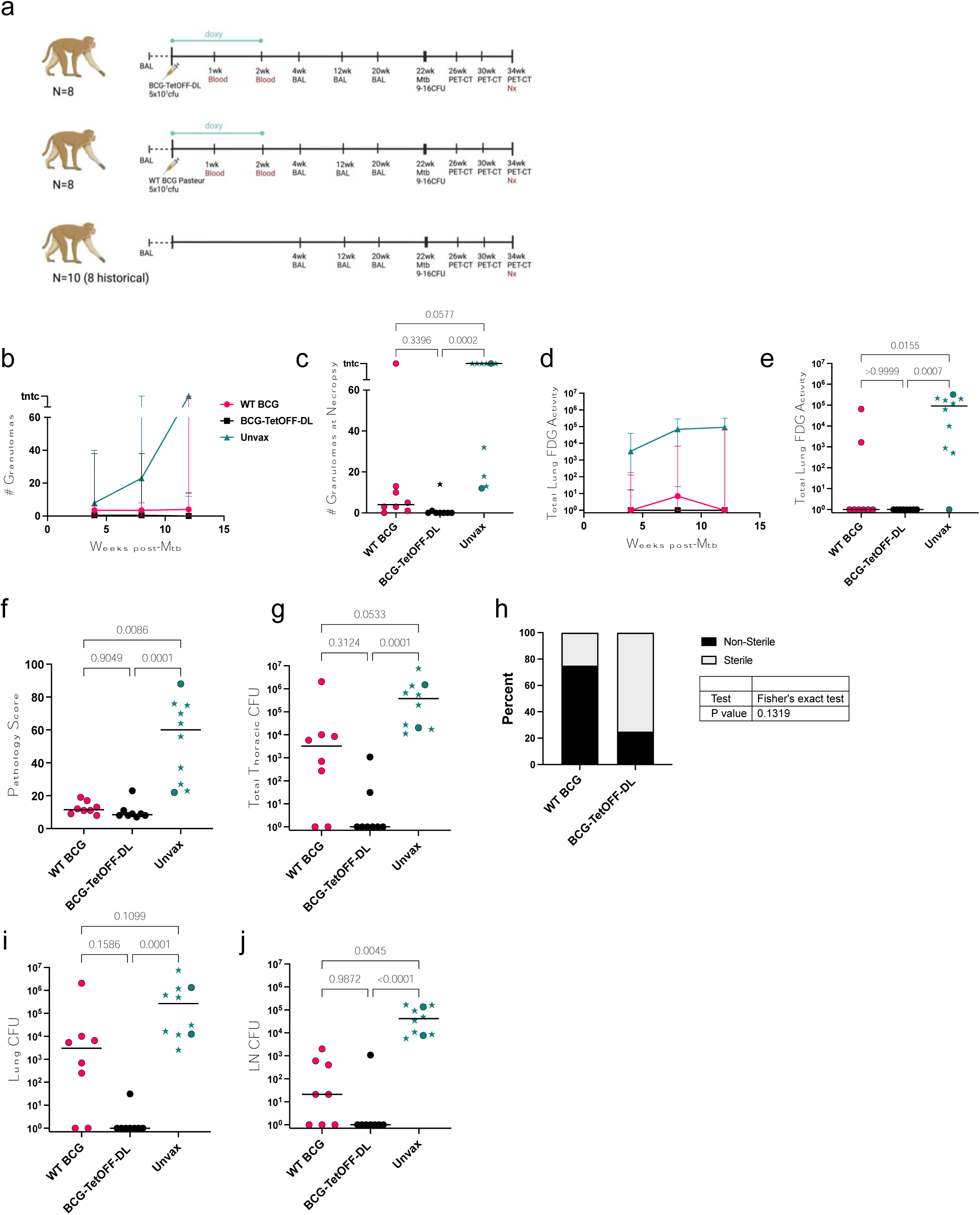
BCG-TetOFF-DL provides robust protection against Mtb infection in macaques. (a) Efficacy study design. (b, c) Number of lung granulomas over time (b) and at necropsy (c). (d, e) Total lung FDG activity by PET imaging over time (d) and at necropsy (e). (f) Gross pathology scores at necropsy. (g) Total thoracic bacterial burden at necropsy. (h) Fisher’s exact test showing trend of higher levels of sterility in animals vaccinated with BCG-TetOFF-DL versus WT BCG Pasteur; Fisher’s exact p-value reported. Total bacterial burden in lungs (i) and thoracic lymph nodes (j). WT BCG (n=8), BCG-TetOFF-DL (n=8) and unvaccinated animals (n=10). Stars represent historical controls. Lines represent the median and range. Statistic for c, e, f, g, i, j Kruskal-Wallis tests were performed and Dunn’s p-values (adjusted for multiple comparisons) were reported.

### T-cell responses in airways were similar upon vaccination with BCG-TetOFF-DL or WT BCG Pasteur

We monitored the immune response in the airways for both vaccine strains and unvaccinated control animals using BALs. BALs were performed pre-vaccination and 4, 12 and 20 weeks post IV-vaccination. We assessed the number of memory effector T-cells at each time point (memory CD4+ or memory CD8+ producing either IFN- γ, TNF, IL-17, IL-2, GzmB, GzmK, granulysin or perforin) (**Fig. S6**). Both WT and BCG-TetOFF-DL induced a sustained increase in total T cell numbers in BAL while there was little change in the unvaccinated animals. This corresponded to a sustained increase in T cells producing cytokines and in CD8 T cells producing perforin in vaccinated animals. This confirms and extends our data from the persistence study (**Fig. 4**) that a robust T cell response is generated upon vaccination with both IV BCG-TetOFF-DL and WT BCG Pasteur. At 20 weeks, the final time point before challenge, we did not observe significant differences in effector T-cell numbers in the airway when comparing BCG-TetOFF-DL and WT BCG Pasteur. The similar level of response in both vaccination groups suggests live mycobacteria are only required to be present for a short amount of time for a robust memory response to be generated.

### BCG-TetOFF-DL and WT BCG had fewer granulomas and less lung inflammation compared to unvaccinated macaques

Serial PET-CT allows the assessment of disease trajectory by enumerating numbers of granulomas and lung and lymph node inflammation over time; increasing numbers of granulomas indicates Mtb dissemination which correlates with development of active TB. Thus, greater the granuloma count, the more severe the infection and the less protective the vaccine. NHPs vaccinated with either BCG-TetOFF-DL or WT BCG Pasteur exhibited lower granuloma formation at every time point compared to unvaccinated control animals (**Fig. 5b,c**). One WT BCG Pasteur vaccinated NHP developed a large number of granulomas. Higher levels of the PET probe ^18^F fluorodeoxyglucose (FDG) activity in the lung due to increased host cell glucose metabolism indicate an increase in lung inflammation ^21,22^. We previously demonstrated that total lung FDG activity is correlated to bacterial burden in Mtb infected macaques ^2^. Unvaccinated animals had significantly more total lung FDG activity after Mtb challenge compared to vaccinated animals (**Fig. 5d,e**). All animals of the BCG-TetOFF-DL vaccinated group had undetectable levels of inflammation (via lung FDG) just prior to necropsy, whereas 2 of the 8 animals in the WT BCG Pasteur group had elevated levels of total lung FDG activity (**Fig. 5e**).

### BCG-TetOFF-DL vaccination leads to enhanced protection against Mtb challenge

At necropsy, gross pathology scores in vaccinated animals were significantly lower than unvaccinated animals (**Fig. 5f**). Our stated primary outcome measure of vaccine efficacy was total thoracic Mtb burden at necropsy. BCG-TetOFF-DL IV vaccinated animals displayed robust protection against Mtb with significantly lower total thoracic CFU compared to unvaccinated animals (p=0.001)(**Fig. 5g**). WT BCG IV vaccinated animals also had lower bacterial burdens compared to unvaccinated macaques (p=.0583)(**Fig. 5g**). Although BCG-TetOFF-DL was not significantly different than WT BCG in terms of total thoracic bacterial burden (p=0.3124), 6 out of the 8 animals in the IV-BCG-TetOFF-DL were sterile, defined as 0 Mtb CFU recovered, compared to two of eight for WT BCG (**Fig. 5h**). Thus, there was a trend towards increased sterile protection in the BCG-TetOFF-DL macaques (Fisher’s exact test, p = 0.1319). Total thoracic bacterial burden can be separated into lung CFU and thoracic lymph node (LN) CFU. BCG-TetOFF-DL vaccinated animals had significantly reduced lung CFU, with only one of the eight animals with lung CFU, while there was only a trend towards lower lung CFU in the WT BCG animals compared to unvaccinated animals (**Fig. 5i**). Both BCG-TetOFF-DL and WT BCG IV vaccinated animals had significantly lower thoracic LN CFU compared to unvaccinated animals (**Fig. 5j**), suggesting dissemination from lung to lymph nodes was reduced with both WT and BCG-TetOFF-DL.

### CD4 T cell responses are enhanced in the lungs of BCG-TetOFF-DL compared to WT BCG vaccinated macaques

The bacterial burden data suggest better protection by BCG-TetOFF-DL IV vaccination than by WT BCG using the Pasteur strain (6/8 sterile with BCG-TetOFF-DL vs 2/8 for WT BCG). To investigate the factors that might contribute to this, we analyzed immune cell populations and functions in the tissues of the macaques at necropsy. Multiparametric spectral flow cytometry and Boolean gating revealed a significantly higher frequency of CD4+ T cells in lungs of BCG-TetOFF-DL animals, whereas the CD8+ T cell population was slightly, although not significantly, higher in WT BCG Pasteur vaccinated animals (**Fig. 6a**). There was a significant increase in the frequency of lung memory CD4+ T-cells producing cytokines (IFN-γ, TNF, IL-17 and/or IL-2) in the BCG-TetOFF- DL vaccinated group compared to the WT BCG Pasteur group upon stimulation with mycobacterial WCL (**Fig. 6b**). There were no statistically significant differences between BCG-TetOFF-DL or WT BCG vaccinated animals in lung T cells producing cytotoxic effector molecules, although slightly higher frequencies were seen in WT BCG vaccinated animals (**Fig. 6b**). In the thoracic lymph nodes, WT BCG vaccinated animals had significantly higher frequencies of CD8+ T cells producing cytotoxic molecules compared to BCG-TetOFF-DL vaccinated animals (**Fig. 6d**). There was an increase in the number of cytokine producing memory CD4+ cells, specifically IFN-γ, IL- 2 and TNF, present in the lung tissue of BCG-TetOFF-DL vaccinated macaques compared to those vaccinated with WT BCG (**Fig. S7**), corroborating the significant increase in frequency of CD4 T cells.

**Figure 6.**
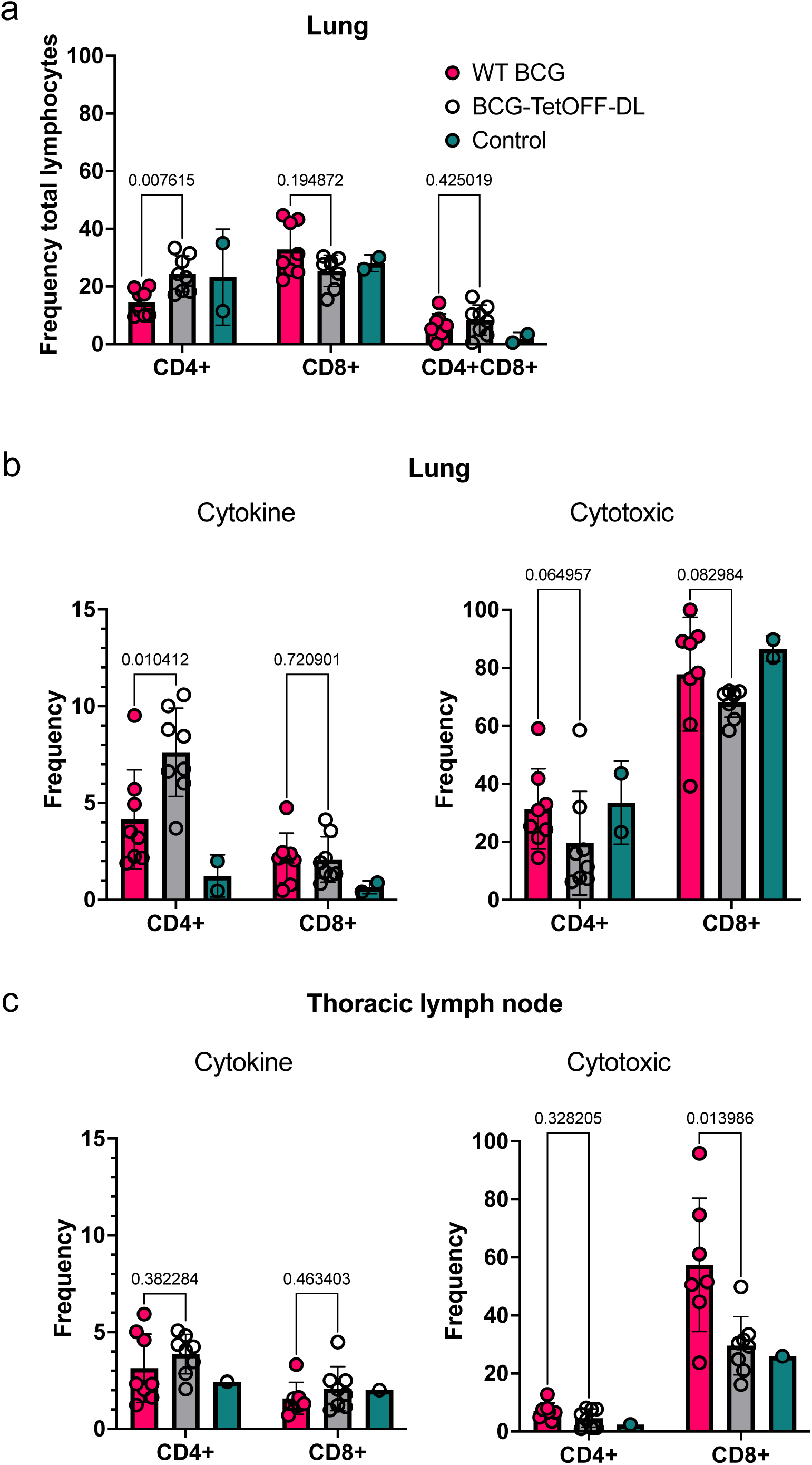
BCG-TetOFF-DL induces more CD4 T cells producing cytokines in lung tissue compared to WT BCG. (a) Frequency of CD4+ and CD8+ T-cells as a percentage of CD3+ cells isolated from homogenized lung tissue at necropsy (WT BCG, BCG-TetOFF-DL n=8, unvaccinated n = 2). Frequency of effector CD4+ and CD8+ T-cells in the lung (b) and thoracic lymph nodes (c) producing cytokines or cytotoxic molecules. Mean with SD shown. Statistics: Mann-Whitney p values reported comparing BCG-TetOFF-DL and WT BCG.

### Spleen pathology

Due to the systemic nature of IV BCG vaccination and the robust immune response induced, splenomegaly was observed in NHPs ^2^. The duration of the enlargement and the relationship with BCG survival is unknown. Here we show that spleen size appears to correlate with time post-vaccination and BCG CFU (**Fig. S**8). IV-BCG vaccinated and IV-BCG vaccinated/Mtb challenged animals had spleen sizes that were larger than the average range of an unvaccinated/unchallenged macaque but were largest at 4 weeks post vaccination. Spleen size reduced over time, with the smallest measurement being recorded 34 weeks post vaccination. Even spleens from unvaccinated but Mtb challenged macaques did not fall within the normal range. We assume that this is due to Mtb infection in these animals, noting that IV BCG vaccinated and Mtb challenged animals had spleen sizes similar to unvaccinated and challenged macaques at this late time point.

## Discussion

BCG has likely been administered to more humans than any other vaccine designed to prevent infectious diseases and is generally safe ^23^. Though rare (0.1 to 4.3 per one million vaccinated children), complications from, BCG vaccination is one of the most common causes of death in immunocompromised children ^24^. Although BCG is an effective option for treatment of bladder cancer, approximately 8% of patients develop complications which leads to cessation of treatment ^11^. We therefore sought to generate BCG strains whose elimination does not depend on a patient’s immune system. Usage of phage lysins to establish conditional kill switches was prioritized over other toxins because we expected these enzymes to not only increase safety but to also increase immunogenicity via the release of cytoplasmic antigens.

Combining the controlled expression of two phage lysins resulted in BCG strains that either require atc/doxy to grow or are efficiently killed by exposure to atc/doxy and escape from this controlled growth was as low as ∼2 x 10^-9^ mutants per cell division. Escape was thus not as low as for the Mtb strain described in the accompanying manuscript (Wang et. al. submitted) but likely low enough to allow studies in humans given that BCG is attenuated already. Characterization of the atc/doxy-addicted BCG (BCG-TetOFF-DL) in SCID mice confirmed that induction of lysin expression is sufficient to eliminate the strain from the lungs and reduce bacterial burden in the spleens in the absence of an intact immune system. Mice vaccinated with BCG-TetOFF-DL or WT BCG showed similar protection against Mtb challenge, even though the BCG-TetOFF-DL strain was eliminated faster than WT BCG. It seems likely that protection benefited from lysis of BCG-TetOFF-DL as it releases additional antigens. This interpretation is supported by the enhanced stimulation of proinflammatory cytokines we observed in macrophages infected with this strain compared to WT BCG.

BCG delivered intravenously was shown to provide robust protection against Mtb infection and disease in macaques ^2,25^. Here we show, using genetically modified BCG that is killed by cessation of doxy administration in vivo in macaques, that BCG does not need to be alive for more than a few weeks to provide robust protection against Mtb challenge. Our data indicate that this “suicide” BCG strain induces greater CD4 T cell responses in lungs and may provide even more robust protection than WT BCG in macaques. Using a self-killing BCG strain may thus increase the safety of IV BCG vaccination strategies while maintaining remarkable protective efficacy.

The ability of BCG-TetOFF-DL to generate a robust CD4 response could be vital in its ability to be an efficacious vaccine, as CD4 T cells are known to play an important role in protection against TB ^26^. Our data indicated that BCG-TetOFF-DL induces a stronger CD4 T cell response compared to WT BCG with production of key cytokines including IFN-γ and TNF in lungs. We hypothesize that the lysis of BCG-TetOFF-DL leading to release of mycobacterial proteins and the enhanced cytokine production by macrophages, as shown in vitro, leads to a more robust priming of T cells in vivo, at least in macaques.

BCG IV was previously noted to result in enlarged spleens in macaques. We considered that reducing the duration of live BCG could mitigate this side effect. In the persistence study, at 4 weeks post-BCG-TetOFF-DL, spleens were quite large, but reduced in size by 8 weeks. Spleen sizes were smaller overall in the BCG-TetOFF- DL vaccinated macaques that received doxy for only 2 weeks compared to those receiving doxy for 8 weeks. In the Mtb challenged animals, both vaccinated and unvaccinated animals had increased spleen sizes compared to “normal” spleens and were slightly smaller than the spleens harvested at 8 weeks post-vaccination, although spleen size can vary with age, sex, and size of macaques. Mtb infection generates immune responses resulting in spleen enlargement, which may suggest the normal spleen size range is not achievable in Mtb infected groups^27^. The BCG associated non-necrotizing ‘microgranulomas’ that were present 4- and 8-weeks post-vaccination were not seen in animals necropsied at 34 weeks post-vaccination.

BCG-TetOFF-DL was generated using a BCG Pasteur background and the WT BCG used as a comparator here was the same Pasteur strain, while our previous IV-BCG studies performed in NHPs used BCG-SSI (Danish) as the vaccine strain ^2,25,28^. Thus, both BCG Pasteur and BCG Danish/SSI can achieve sterilizing levels of protection when delivered IV. This study and one other BCG IV study ^28^ were performed in Mauritian cynomolgus macaques (MCM) while our original study was performed in rhesus macaques. MCM have similar susceptibility to TB disease as rhesus macaques ^21,29^ and BCG IV provides robust protection in both macaque ^2,25,28^. In contrast, BCG-TetOFF-DL or WT BCG delivered IV provided only modest protection in mice in the current study, similar to other studies with IV BCG in mice ^30,31^. Thus, vaccines that provide robust protection in macaques cannot always be predicted from murine data.

Limitations to this study are the small samples sizes used in the macaque persistence study which limited statistical analyses. Similarly, larger sample sizes for the macaque protective efficacy study could provide clearer differences between BCG-TetOFF-DL and WT BCG outcomes. Future studies could also assess whether doxy is necessary in vivo, or whether the strain could be eliminated even earlier in the vaccination phase. In mice it remains to be demonstrated whether vaccination efficacy can be improved by prolonging persistence of the BCG kill switch strain by administrating doxy for different periods of time.

The safety and efficacy of IV-BCG in humans is yet to be determined and there are practical challenges to implementing BCG IV for widespread vaccination. One concern is the increased spleen sizes in macaques following BCG IV and whether the systemic spread of live-attenuated bacteria may result in human illness. The “suicide” strain developed in this study aimed to resolve some of these issues, while maintaining the high levels of protection seen with WT IV-BCG vaccination. Although the splenomegaly was still present, spleen size did reduce over time. We did not observe any adverse effects or signs of illness in the macaques following WT or BCG-TetOFF-DL vaccination, and blood cultures during the early phase of vaccination were negative. In our previous study in SIV+ macaques given WT BCG, blood cultures were also negative 2 weeks post-vaccination

In summary, our data support that a limited exposure to BCG delivered intravenously is effective against Mtb challenge. The use of a strain that lyses itself may provide enhanced protection due to increased immunogenicity in vivo. This “suicide” BCG strain could limit safety concerns that are raised regarding IV administration of a live vaccine as well as provide an option for safer intradermal BCG vaccination of immunocompromised individuals and treatment for bladder cancer.

## Materials and Methods

### Strains, media and culture conditions

All *M. bovis* BCG strains are derived from BCG Pasteur TMC 1011 obtained from the American Type Culture Collection (ATCC #35734). *M. tuberculosis* H37Rv was used for challenge experiments. Strains were cultured in liquid Middlebrook 7H9 medium supplemented with 0.2% glycerol, 0.05% tween80 and ADN (0.5% bovine serum albumin, 0.2% dextrose, 0.085% NaCl) and on Middlebrook 7H10 agar plates supplemented with 0.5% glycerol and Middlebrook OADC enrichment (Becton Dickinson). Antibiotics were added for selection of genetically modified strains at the following concentrations: hygromycin (50 μg/ml), kanamycin (25 μg/ml), zeocin (25 μg/ml). Anhydrotetracycline was used at 0.5 or 1 μg/ml.

### Generation of strains

To create the TetON single and dual-lysin strains, BCG was transformed with either or both plasmids pGMCK3-TSC10M1-D29L and pGMCgZni-TSC10M1-L5L, integrating the D29-lysin and L5-lysin into BCG L5 and Giles sites, respectively. To create the TetOFF single and dual-lysin strains, BCG was transformed with either or both plasmids pGMCK3-TSC10M-TsynC-pipR-SDn-P1-TsynE-PptR-L5L and pGMCgZni-TSC38S38-TrrnBd2-P749pld-10C32C8C-D29L, integrating the L5-lysin and D29-lysin into BCG L5 and Giles sites, respectively. The single lysin strains were cultured in the presence of 0.5 μg/ml atc and the dual lysin strain was cultured in the presence of 1 μg/ml atc.

### Analysis of bacterial lysis by immunoblotting

Culture filtrates were prepared as follows. BCG strains were grown in 7H9 medium with 0.2% glycerol, 0.05% Tween-80, 0.5% BSA, 0.2% dextrose and 0.085% NaCl until the culture reached an OD of 0.6 ∼ 0.8. Cultures were then washed three times with PBS to remove BSA and Tween-80. We next suspended the pellet in 7H9 medium supplemented with 0.2% glycerol, 0.2% dextrose and 0.085% NaCl. After incubation, culture supernatant was harvested by centrifugation and filtration through 0.22 μm filters. Filtrates were concentrated 100-fold by using 3K centrifugal filter units (Millipore) and analyzed by immunoblotting with antisera against Eno and PrcB.

### Fluctuation analysis

Fluctuation analysis was performed as previously reported ^32^. Briefly, BCG strains were inoculated at permissive condition (1.0 µg/mL atc) in 7H9 media supplemented with OADC in the presence of antibiotics (20 µg/mL kanamycin, 25 µg/mL zeocin). After reaching OD=1.0, the culture was diluted to multiple 4-mL aliquots with 1,000 bacteria. The diluted culture was grown for 11-14 days in 7H9+OADC media in permissive conditions in the presence of antibiotics. Once OD was at 1.0, bacteria were washed for three times and resuspended in 400 µL 7H9+OADC without atc. Four aliquots of bacteria were streaked onto 7H10+OADC plates supplemented with antibiotics (20 µg/mL kanamycin, 25 µg/mL zeocin) and 1.0 µg/mL atc for bacteria count, and the rest aliquots were spread onto 7H10+OADC plates with antibiotics (20 µg/mL kanamycin, 25 µg/mL zeocin) and either atc, or no atc. According to Ma, Sarkar, Sandri (MSS) method ^33^, the estimated number of mutations per culture (*m*) was inferred by number of mutant (*r*) colonies observed on plates. The escape rate was calculated by dividing *m* by *N_t_*, the number of cells plated for each culture. The Mann-Whitney *U* test was used to statistically compare escape rates between two groups. The lowest detection limit was calculated based on an assumption that only one colony could be observed in all 20 independent cultures.

### Preparation and infection of murine bone marrow derived macrophages (BMDMs)

Femurs and tibias of female C57BL/6 mice were extracted, and bone marrow cells were aseptically flushed using PBS. Cells were re- suspended in Dulbecco’s modified eagle medium (DMEM) supplemented with 10% fetal bovine serum (FBS) and 20% L929 culture filtrate and incubated for 6 days to allow differentiation into macrophages. Cells were harvested and seeded at 6 x 10^4^ per well in 96-well plates in DMEM supplemented with 10% FBS, glycine and 10% L929 cell culture overnight before infection. Mycobacteria were washed in PBS + 0.05% tyloxapol and a single cell suspension was generated by low-speed centrifugation to pellet clumped cells. The bacteria were diluted into DMEM, 10% LCM and added to macrophages at MOI of 0.1. After 4 hours, extracellular bacteria were removed by washing the macrophages three times with warm PBS. Infected BMDMs were treated with atc (0.5 μg/ml) or RIF (0.5 μg/ml) starting 16 hrs post infection. Cytokines were quantified using BD OptEIA ELISA kits for mouse TNF, IL-12p40 or IL6 (BD Biosciences). The number of intracellular bacteria was determined by lysing macrophages with 0.01% Triton X-100 and culturing dilutions of macrophage lysates on 7H10 agar plates.

### Mouse infections

Female 8- to 10-week-old C57BL/6 (# 000664, Jackson Laboratory) were vaccinated with ∼ 10^6^ CFU of the indicated BCG strain by the intravenous route. Mice received doxycycline containing mouse chow (2,000 ppm; Research Diets) for the indicated periods. Mice were infected with Mtb H37Rv using an inhalation exposure system (Glas-Col) with a mid-log phase Mtb culture to deliver approximately 100 bacilli per mouse. To enumerate CFU, organs were homogenized in PBS and cultured on 7H10 agar. Charcoal (0.4 %, w/v) was added to the plates that were used to culture homogenates from doxy treated mice. Agar plates were incubated for 3- 4 weeks at 37°C. Mice were housed in a BSL3 vivarium. All mouse experiments were approved by and performed in accordance with requirements of the Weill Cornell Medicine Institutional Animal Care and Use Committee.

### Flow cytometry to assess immune responses in mice

Mouse lungs were isolated and placed in RPMI1640 containing Liberase Blendzyme 3 (70 μg/ml; Roche) and DNase I (50 μg/ml; Sigma-Aldrich). Lungs were then cut into small pieces and incubated at 37 C for 1 hour. The cells were filtered using cell strainers, collected by centrifugation, resuspended in ACK hemolysis buffer (ThermoFisher) and incubated for 10 minutes at room temperature. Cells were then washed with PBS and resuspended in splenocyte medium (RPMI-1640, supplemented with 10% FBS, 2 mM GlutaMax 10 mM HEPES, and 50 μM 2-mercaptoethanol). For intracellular staining samples, cells were stimulated with PPD (20 μg/ml) in the presence of anti-CD28 antibody (37.51, BioLegend) for 1.5 hours and 10 μg/ml Brefeldin A (Sigma) and monesin were added and incubated at 37 C for another 3 hrs. Samples were kept on ice in a refrigerator overnight. Cells were stained with Zombie-NIR (BioLegend) to discriminate live and dead cells. Purified anti-CD16/32 antibody (93, BioLegend) was used to block Fc receptor before staining. PerCp-Cy5.5 anti-CD62L (MEL-14, Thermofisher), BV605 anti-CD4 (RM4-5, BioLegend), BV711 anti-CD8 (53-6.7, BioLegend), eFluor450 anti-CD11a (M17/4, Thermofisher), BUV395 anti- CD153 (RM153, BD Biosciences), BV480 anti-CD69 (H1.2F3, BD Biosciences), APCC7 anti-CD44 (IM7, Biolegend) were used to stain cells for 30 minutes at room temperature. The cells were fixed in fixation buffer (BioLegend) for 30 minutes and taken out of BSL3. The cells were incubated for 20 minutes in permeabilization buffer (eBioscience) before intracellular cytokine staining. BV421 anti IL-17A (TC11-18H10.1, Biolegend), BV750 anti-IFNγ (XMG1.2, BD Biosciences), PE-C5.5 anti-IL-2 (JES6-5H4, Biolegend), FITC anti-TNF (MP6-XT22, BioLegend), BV785 anti-CD3 (17A2, BioLegend) were used to stain cells for 30 minutes. Cells were washed and resuspend in cell staining buffer (BioLegend). Flow cytometry data were acquired on cytometer (LSR Fortessa TM; BD Biosciences) or cytometer (Cytek Aurora; Cytek Biosciences) and were analyzed with FlowJoTM V10.

### Macaques

Mauritian cynomolgus macaques (*Maccaca fasicularis*) used in this study were obtained from BioCulture US (all males, 6-9 years old). All procedures and study design complied with ethical regulations and were approved by the Institutional Animal Care and Use Committee of the University of Pittsburgh. Macaques were housed either singly or in pairs in a BSL2 animal facility and cared for in accordance with local, state, federal, and institute policies in facilities accredited by the American Association for Accreditation of Laboratory Animal Care (AAALAC), under standards established in the Animal Welfare Act and the Guide for the Care and Use of Laboratory Animals. During the Mtb challenge phase, animals were housed in a BSL3 animal facility. Macaques were monitored for physical health, food consumption, body weight, temperature, complete blood counts, and serum chemistries. Full details on macaques in this study are in **Table S3**.

### BCG vaccination

To assess persistence of BCG-TetOFF-DL, animals that were intravenously vaccinated were sedated with ketamine (10 mg/kg) or telazol (5- 8 mg/kg) and injected intravenously with 3.74 x 10^7^ CFU BCG-TetOFF-DL vaccine in an injection volume of 1mL. Animals that underwent challenge with *M. tuberculosis* strain Erdman were vaccinated intravenously with 5 x 10^7^ CFU BCG-TetOFF-DL or WT BCG Pasteur.

### Macaque Mtb challenge

Macaques were challenged by bronchoscope with 9-16 Mtb strain Erdman 22 weeks post vaccination as previously described ^34^. Control animals (unvaccinated) were challenged at the same time. Historical control unvaccinated and Mtb challenged MCMs (previously published by our group ^28^) were included for statistical analyses and comparison.

### Blood, BAL and tissue processing

To assess the presence of viable BCG-TetOFF-DL, blood was drawn from each animal 1- and 2-weeks post vaccination. Blood was cultured and the presence of BCG-TetOFF-DL assessed using atc-containing plates. Bronchoalveolar lavages for the persistence study in macaques were performed pre-vaccination, 4 weeks post vaccination (all groups) and 8 weeks post vaccination (for groups B and C). For the immunization and challenge studies in macaques, BAL was performed prior to vaccination and monthly thereafter until the time of challenge. Procedures were performed as previously described ^2^. PBMCs were isolated from peripheral blood using Ficoll-Paque PLUS gradient separation (GE Healthcare Biosciences) and standard procedures.

### PET/CT

PET/CT scans were performed at designated time points throughout the study to assess thoracic cavity inflammation. Animals were sedated with 10 mg/kg ketamine / 0.5 ml atropine before imaging. An intravenous catheter was placed in the saphenous vein and animals were injected with ∼ 5 millicurie (mCi) of ^18^F-FDG. Once placed on the imaging bed, anesthesia was induced with 2.5 to 3% isoflurane which is reduced to 0.8–1.2% for maintenance. Breathing during imaging was maintained using an Inspiration 7i ventilator (eVent Medical, Lake Forest, CA, USA) with the following settings: PF = 9.0 l/min, respiration rate = 18–22 bpm, tidal volume = 60 ml, O2 = 100, PEEP = 5–8 cm H_2_O, peak pressure = 15–18 cm H_2_O, I:E ratio = 1:2.0. A breath hold was conducted during the entirety of the CT acquisition.

PET/CT scans were performed on a MultiScan LFER 150 (Mediso Medical Imaging Systems, Budapest, Hungary). CT acquisition was performed using the following parameters: Semi-circular single field-of-view, 360 projections, 80 kVp, 670 μA, exposure time 90 ms, binning 1:4, voxel size of final image: 500 x 500 μm. PET acquisition was performed 55 min after intravenous injection of ^18^F-FDG with the following parameters: 10 min acquisition, single field-of-view, 1–9 coincidence mode, 5 ns coincidence time window. PET images were reconstructed with the following parameters: Tera-Tomo 3D reconstruction, 400–600 keV energy window, 1–9 coincidence mode, median filter on, spike filter on, voxel size 0.7 mm, 8 iterations, 9 subsets, scatter correction on, attenuation correction based on CT material map segmentation. Serial CT or PET/CT images were acquired pre-infection and at 4 and 11 dpi. Animal A2 was CT scanned at 9 dpi instead of 11 dpi, and animal A1 was scanned at 18 dpi in addition to the standard imaging schedule previously described.

Images were analyzed using OsiriX MD or 64-bit (v.11, Pixmeo, Geneva, Switzerland). Before analysis, PET images were Gaussian smoothed in OsiriX and smoothing was applied to raw data with a 3 x 3 matrix size and a matrix normalization value of 24. Whole lung FDG uptake was measured by first creating a whole lung region- of-interest (ROI) on the lung in the CT scan by creating a 3D growing region highlighting every voxel in the lungs between -1024 and -500 Hounsfield units. This whole lung ROI was copied and pasted to the PET scan and gaps within the ROI were filled in using a closing ROI brush tool with a structuring element radius of 3. All voxels within the lung ROI with a standard uptake value (SUV) below 1.5 were set to zero and the SUVs of the remaining voxels were summed for a total lung FDG uptake (total inflammation) value. Thoracic lymph nodes were analyzed by measuring the maximum SUV within each lymph node using an oval drawing tool. Both total FDG uptake and lymph node uptake values were normalized to back muscle FDG uptake that was measured by drawing cylinder ROIs on the back muscles adjacent to the spine at the same axial level as the carina (SUVCMR; cylinder-muscle- ratio). PET quantification values were organized in Microsoft Excel and graphed using GraphPad Prism.

### Necropsy, pathology scoring, BCG burden and Mtb burden

At necropsy, NHPs were sedated with ketamine and had a maximal blood draw then euthanized by sodium pentobarbital injection, followed by gross examination for pathology. A gross pathology scoring system was employed, assessing lung, lymph node and extrapulmonary compartments ^21^. Spleen size was also measured. Average spleen size of adult macaques was provided by the Wisconsin Primate Center. A pre-necropsy PET-CT scan was used to map lesions in the thoracic cavity and at necropsy these regions were excised and homogenized to form a single-cell suspension. Uninvolved lung tissue, lymph nodes and spleen were also processed. To recover BCG-TetOFF-DL, the individual cell suspensions were plated on 7H11 agar + atc and incubated at 37°C with 5% CO_2_ for 3 weeks with atc replenished on the plates after 10 days of incubation. CFU were counted to assess the BCG-TetOFF-DL burden of each animal. To assess Mtb burden post-challenge, single-cell suspensions were plated on 7H11 agar and incubated at 37°C with 5% CO_2_ for 3 weeks.

### Multiparameter flow cytometry

Up to one million viable cells isolated from tissue or BAL were stimulated with 20 μg/ml H37Rv Mtb whole cell lysate (WCL) (BEI Resources), 1 μg/ml each of ESAT-6 and CFP-10 peptide pools (provided by Aeras, Rockville, MD) or R10 media for 2 h before adding 10 μg/ml BD GolgiPlug (BD Biosciences) for a further 12 hours. Cells were surface stained to allow for the assessment of cell composition, followed by intracellular staining (ICS) for the analysis of cytokine and cytotoxic molecule production. Permeabilization for ICS was performed using BD Fixation/Permeabilization Kit. Surface and ICS antibody cocktails were made in BD Brilliant Stain buffer. Cells were analyzed using a five laser Cytek® Aurora spectral flow cytometer. The flow cytometry results were analyzed using FlowJo™ v10.8 Software (BD Life Sciences). All antibodies used in the flow panels are shown in **Table S1** and **Table S2**. Gamma/delta (TCR106, Invitrogen) and perforin (3465-3-500, Mabtech) antibodies were conjugated with PE/Cy5® Conjugation Kit (ab102893, abcam) and Alexa Fluor® 488 Conjugation Kit - Lightning-Link® (ab236553, abcam) respectively. Gating strategy is presented in **Fig. S9**.

#### Statistical methods

For mice, generation of graphics and data analyses were performed in Prism version 10.0.2 software (GraphPad). NHP data were tested for normality using the Shapiro-Wilk test. For comparisons between only BCG-TetOFF-DL and WT, Mann-Whitney tests were used. For comparisons including historical unvaccinated macaque controls, Kruskal-Wallis tests were performed and Dunn’s p-values (adjusted for multiple comparisons) were reported. For longitudinal data with only 3 animals per group, two-way repeated measure ANOVA was performed (random variable was NHP) with the assumption of sphericity. For BAL longitudinal data, groups were compared at each time point using unpaired t-tests (with Welch correction) and Holm-Šídák adjustment for multiple comparisons. For categorical variables, Fisher’s exact test p-value was reported. Statistical tests were not run for any groups with 3 or fewer points. All p-values less than 0.10 are shown.

#### Data Availability Statement

All relevant data are available from the corresponding author upon reasonable request.

## Acknowledgements

We thank Eric Rubin, Sarah Fortune, Kristine Guinn at Harvard University and Robert A. Seder, Patricia Darrah and Mario Roederer at the NIH for helpful discussions. We thank Rodrigo Aguilera Olvera, Heather Kim and Anthony Castro for help with the mouse infection studies. We thank Sophie Lavalette-Levy, Anisha Zaveri and Natalie Thornton for their help with cloning and data analysis. We thank Erica Larson and Chuck Scanga for providing historical macaque control data. We are grateful to the veterinary and research technicians in the Flynn lab for their dedication and expertise in conducting these studies. Research reported in this publication was supported by NIH R01 AI143788, by Leidos Biomedical Research Inc and the National Cancer Institute of the NIH under Contract Number HHSN2612015000031 and NIH NIAID Contract 75N93019C00071. The content is solely the responsibility of the authors and does not necessarily represent the official views of Leidos Biomedical Research, Inc. or the National Institutes of Health.

---

**Figure S1. Single and dual lysin kill switches**.

(a) TetON single lysin kill switch schematic. Lysin expression is repressed by tetracycline repressor (TetR) and induced by anhydrotetracycline (atc) or doxycycline (doxy) which bind TetR and prevent it from binding DNA.

(b) Impact of atc on growth of BCG-TetON single lysin strains. The paper disc contains 1 μg of atc.

(c) TetOFF dual lysin kill switch schematic. Reverse TetR (Rev TetR) binds to DNA in complex with atc or doxy to repress D29L. TetR represses PipR and in the presence of atc/doxy PipR is expressed and represses L5L.

(d) Impact of atc on growth of BCG-TetOFF-DL. The paper disc contains 1 μg of atc.

**Figure S2. In vitro characterization of BCG kill switch strains.**

(a) Growth of BCG TetON single and dual lysin kill switch strains with and without atc. OD_580_ data are means from duplicate cultures.

(b) Western blot analysis of culture filtrates from BCG-TetON-DL and BCG-TetOFF-DL strains grown in the absence of detergent. A whole cell lysate of WT BCG serves as control. Eno and PrcB were enriched in culture filtrate of BCG-TetON-DL after 6 and 9 days of growth in the presence of atc and in culture filtrate of BCG- TetOFF-DL after 6 and 9 days of growth in the absence of atc indicating cell lysis.

(c) Expression of two lysins reduces the fraction of escape mutants compared to expression of single lysins.Cultures were grown in the presence of atc. Symbols represent data from 3-6 individual cultures and means ± SD are depicted. Significance was determined by one-way analysis of variance (ANOVA) with Tukey’s adjusted p-values shown; **** *P* < 0.0001, ***, p < 0.001.

(d) Growth of BCG TetOFF single and dual lysin kill switch strains with and without atc. Data are means ± SD from triplicate cultures. Error bars are too small to be seen.

(e) Expression of two lysins reduces the fraction of escape mutants compared to expression of single lysins.

Cultures were grown in the absence of atc. Symbols represent data from 6 individual cultures and means ± SD are depicted. Statistical significance was assessed by one-way ANOVA with Tukey’s adjusted p-values shown; **** *P* < 0.0001, ***, p < 0.001.

**Figure S3. Survival of BCG, BCG-TetOFF-DL and BCG-TetON-DL following intravenous vaccination.**

(a) CFU from lungs and spleens of mice infected with BCG and BCG-TetOFF-DL not receiving doxy.

(b) CFU quantification from lungs and spleens of mice infected with BCG and BCG-TetON-DL treated with doxy. Data are means ± SEM from 4-5 mice per group and time point.

(c) CFU quantification from lungs and spleens of BCG-TetOFF-DL infected SCID mice treated or not with doxy for the indicated times. Data are means ± SD from 4 mice per group and time point. Multiple unpaired t-tests run on log_10_-transformed data at each time point with Holm-Šídák adjusted p-values shown. **** p < 0.0001, ***, p < 0.001, ** p < 0.01, * p < 0.05, # 0.05 < p < 0.10, ns p > 0.10.

(d,e) Lung sections stained with hematoxylin and eosin from SCID mice infected with BCG-TetOFF-DL for 84 days. (d) Mice were treated with doxy from day 1-14. (e) Mice were treated with doxy from day 1- 84. Each section is from an individual mouse and is representative of each lung.

**Figure S4. BCG-TetON-DL and wt BCG provide similar protection against Mtb infection in mice.** Mice were i.v. vaccinated with BCG or BCG-TetON-DL or received PBS.

(a,b) Quantification of effector memory CD4 and CD8 T cells in mouse lungs from mice vaccinated with BCG and BCG-TetON-DL. Symbols are data from individual mice with lines indicating mean ± SD. Two-way ANOVA was performed with Tukey’s adjusted p-values shown for each time point. **** p < 0.0001, ***, p < 0.001, ** p < 0.01, * p < 0.05, ns p > 0.10.

(c) Quantification of cytokine producing antigen specific CD4 T cells from mice vaccinated with BCG and BCG- TetON-DL on day 30 post vaccination. Lung cells were restimulated ex vivo with PPD prior to intracellular cytokine staining. Two-way ANOVA with Tukey’s adjusted p-values shown for each cytokine producing population. For most of the cytokine producing cells, there were no statistically significant differences among the treatment groups. **** p < 0.0001, ***, p < 0.001, ** p < 0.01, * p < 0.05, ns p > 0.10.

(d) Bacterial burden in lungs and spleens of vaccinated and PBS treated mice. Mice were infected with Mtb H37Rv by aerosol 90 days post vaccination. Symbols represent data from individual mice with lines indicating mean ± SEM. Two-way ANOVA performed on log_10_-transformed data with Tukey’s adjusted p-values shown at each time point. **** p < 0.0001, ***, p < 0.001, ** p < 0.01, * p < 0.05, ns p > 0.10.

**Figure S5 Histologic analysis indicates microgranulomas in spleen, liver and lymph nodes 8 weeks post-BCG-TetOFF-DL.**

H&E staining of fixed spleen (a), lymph node (b) and liver (c) tissue. (d) Spleen size at necropsy for macaques in persistence study and macaques vaccinated and challenged with Mtb. Dashed lines represent normal spleen size range of adult male MCMs. (d, e) Lymphocyte composition of lung tissue (e, n=4) and thoracic lymph nodes (f, n=3) from NHP in persistence study at necropsy.

**Figure S6 T cell responses in airways are similar between BCG-TetOFF-DL and WT BCG after vaccination.**

Total number of cells, CD4+, CD8+, effector memory CD4+ and CD8+ cells during the vaccination phase with BCG-TetOFF-DL, WT BCG or unvaccinated macaques. BAL samples were obtained pre-vaccination and 4, 12, 20 weeks post vaccination and stimulated with Mtb WCL. Flow cytometry was performed with intracellular staining for effector molecules IFN-γ, TNF, IL-17, IL-2, GzmB, GzmK, granulysin or perforin (vaccinated groups n=8, unvaccinated n=2). Multiple unpaired t-tests were used to compare groups at each time with Holm-Šídák adjusted p-values (# 0.05 < p < 0.10). Median and IQR shown.

**Figure S7 Individual effector molecules produced by T cells in lungs of vaccinated and challenged NHP.**

Total number of CD4+, CD8+, effector memory CD4+ and CD8αβ+ cells in the lung at necropsy (vaccinated groups n=8, unvaccinated n=2). Cells producing either cytokines or cytotoxic molecules (IFN-γ, TNF, IL-17, IL- 2, GzmB, GzmK, granulysin or perforin) were analyzed. Mann-Whitney p-values reported. Median shown.

**Figure S8 Splenomegaly reduces over time post IV-BCG vaccination**

Spleen size at necropsy for macaques in persistence study and macaques vaccinated and challenged with Mtb. Dashed lines represent normal spleen size range of adult male MCMs.

**Figure S9 Gating strategy for flow cytometry**

**Table S1: Flow cytometry panel for NHP samples in persistence study**

**Table S2: Flow cytometry panel for NHP samples in protection study Table**

**S3 Full details on macaques used in this study**

