## Supplemental Figures for "A “suicide” BCG strain provides enhanced immunogenicity and robust protection against *Mycobacterium tuberculosis* in macaques"

**Figure S5 Histologic analysis indicates microgranulomas in spleen, liver and lymph nodes 8 weeks post- BCG-TetOFF-DL.** H&E staining of fixed spleen (a), lymph node (b) and liver (c) tissue. (d, e) Lymphocyte composition of lung tissue (d, n=4) and thoracic lymph nodes (e, n=3) from NHP in persistence study at necropsy.

**Figure S6 T cell responses in airways are similar between BCG-TetOFF-DL and WT BCG after vaccination.** Total number of cells, CD4+, CD8+, effector memory CD4+ and CD8+ cells during the vaccination phase with BCG-TetOFF-DL, WT BCG or unvaccinated macaques. BAL samples were obtained pre-vaccination and 4, 12, 20 weeks post vaccination and stimulated with Mtb WCL. Flow cytometry was performed with intracellular staining for effector molecules IFN-γ, TNF, IL-17, IL-2, GzmB, GzmK, granulysin or perforin. Multiple unpaired t-tests were used to compare groups at each time point with Holm-Šídák multiple comparison adjusted p-values (# $0.05 < p < 0.10$). Median and IQR shown.

**Figure S7 Individual effector molecules produced by T cells in lungs of vaccinated and challenged NHP.** Total number of CD4+, CD8+, effector memory CD4+ and CD8αβ+ cells in the lung at necropsy (n=8). Cells producing either cytokines or cytotoxic molecules (IFN-γ, TNF, IL-17, IL-2, GzmB, GzmK, granulysin or perforin) were analyzed. Mann-Whitney p-values reported. Median shown.

**Figure S8 Splenomegaly reduces over time post IV-BCG vaccination.** Spleen size at necropsy for macaques in persistence study and macaques vaccinated and challenged with Mtb. Dashed lines represent normal spleen size range of adult male MCMs.

**Figure S9 Gating strategy**

**Table S1: Flow cytometry panel for NHP samples**

**Table S2 Full details on macaques used in this study**

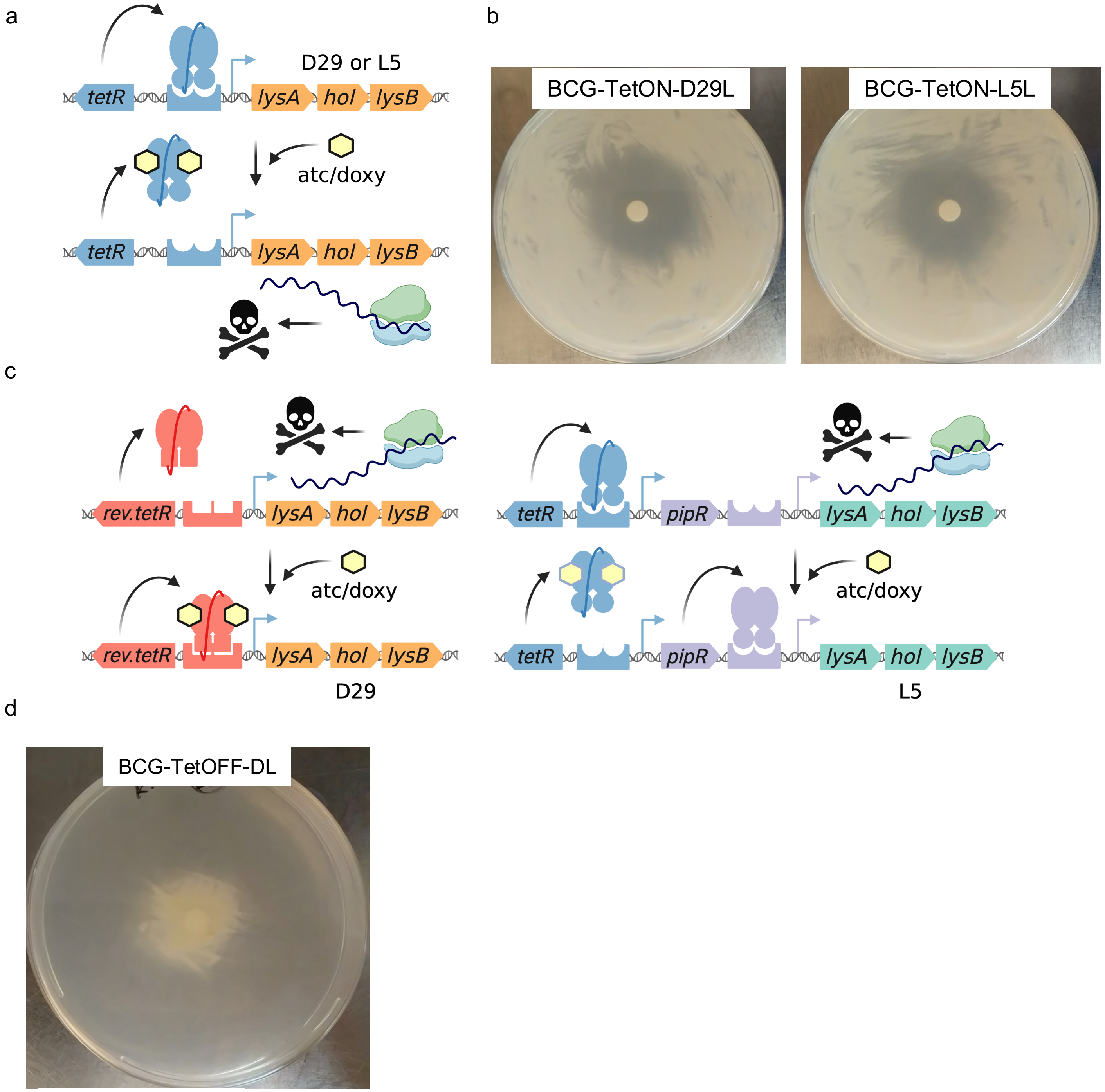
a
b
D29 or L5
tetR
lysA
hol
lysB
atc/doxy
tetR
lysA
hol
lysB
BCG-TetON-D29L
BCG-TetON-L5L
c
rev.tetR
lysA
hol
lysB
atc/doxy
rev.tetR
lysA
hol
lysB
D29
tetR
pipR
lysA
hol
lysB
atc/doxy
tetR
pipR
lysA
hol
lysB
L5
d
BCG-TetOFF-DL

**Figure S1**

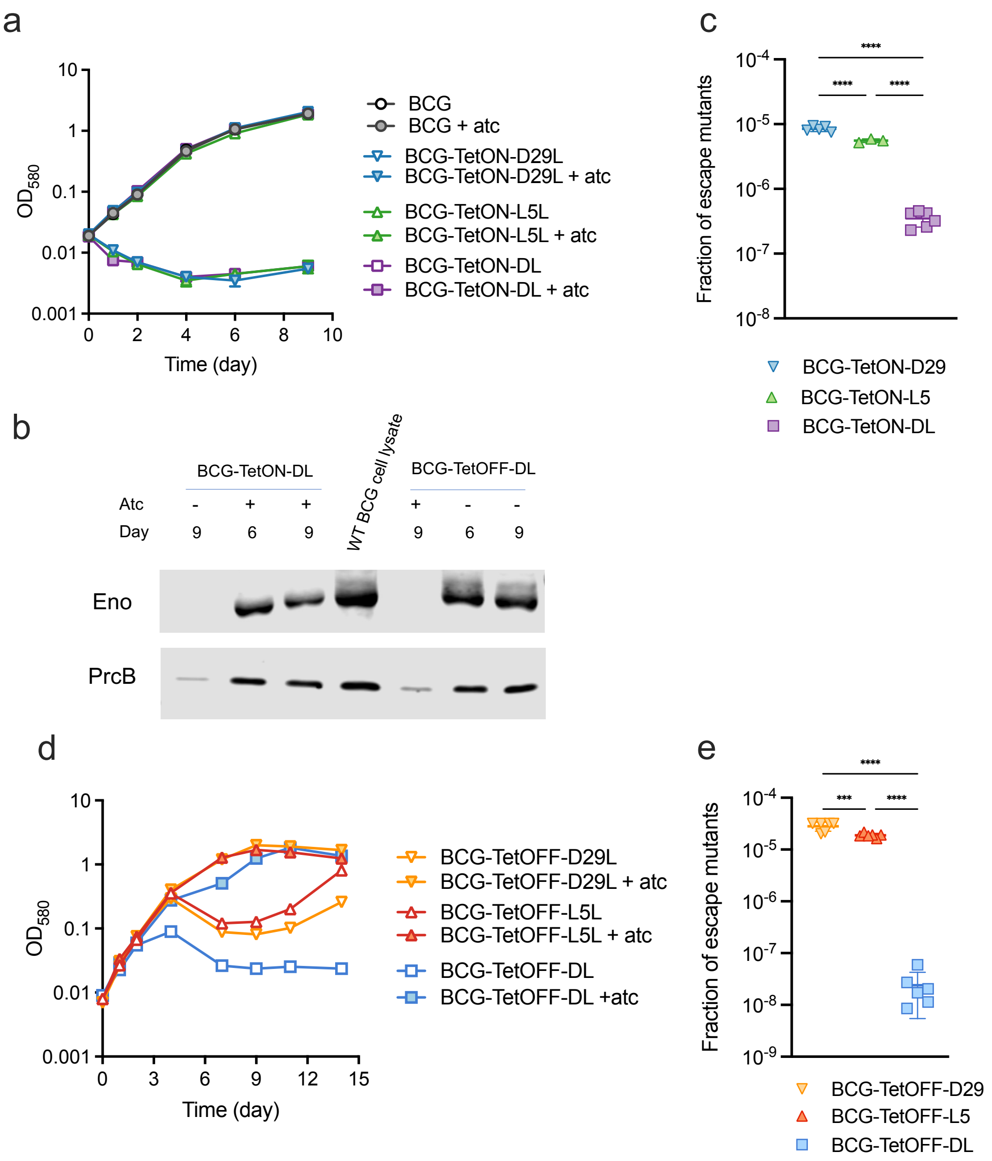
a
OD580
Time (day)
BCG
BCG + atc
BCG-TetON-D29L
BCG-TetON-D29L + atc
BCG-TetON-L5L
BCG-TetON-L5L + atc
BCG-TetON-DL
BCG-TetON-DL + atc
b
BCG-TetON-DL
BCG-TetOFF-DL
WT BCG cell lysate
Atc
Day
Eno
PrcB
c
Fraction of escape mutants
BCG-TetON-D29
BCG-TetON-L5
BCG-TetON-DL
d
OD580
Time (day)
BCG-TetOFF-D29L
BCG-TetOFF-D29L + atc
BCG-TetOFF-L5L
BCG-TetOFF-L5L + atc
BCG-TetOFF-DL
BCG-TetOFF-DL +atc
e
Fraction of escape mutants
BCG-TetOFF-D29
BCG-TetOFF-L5
BCG-TetOFF-DL

**Figure S2**

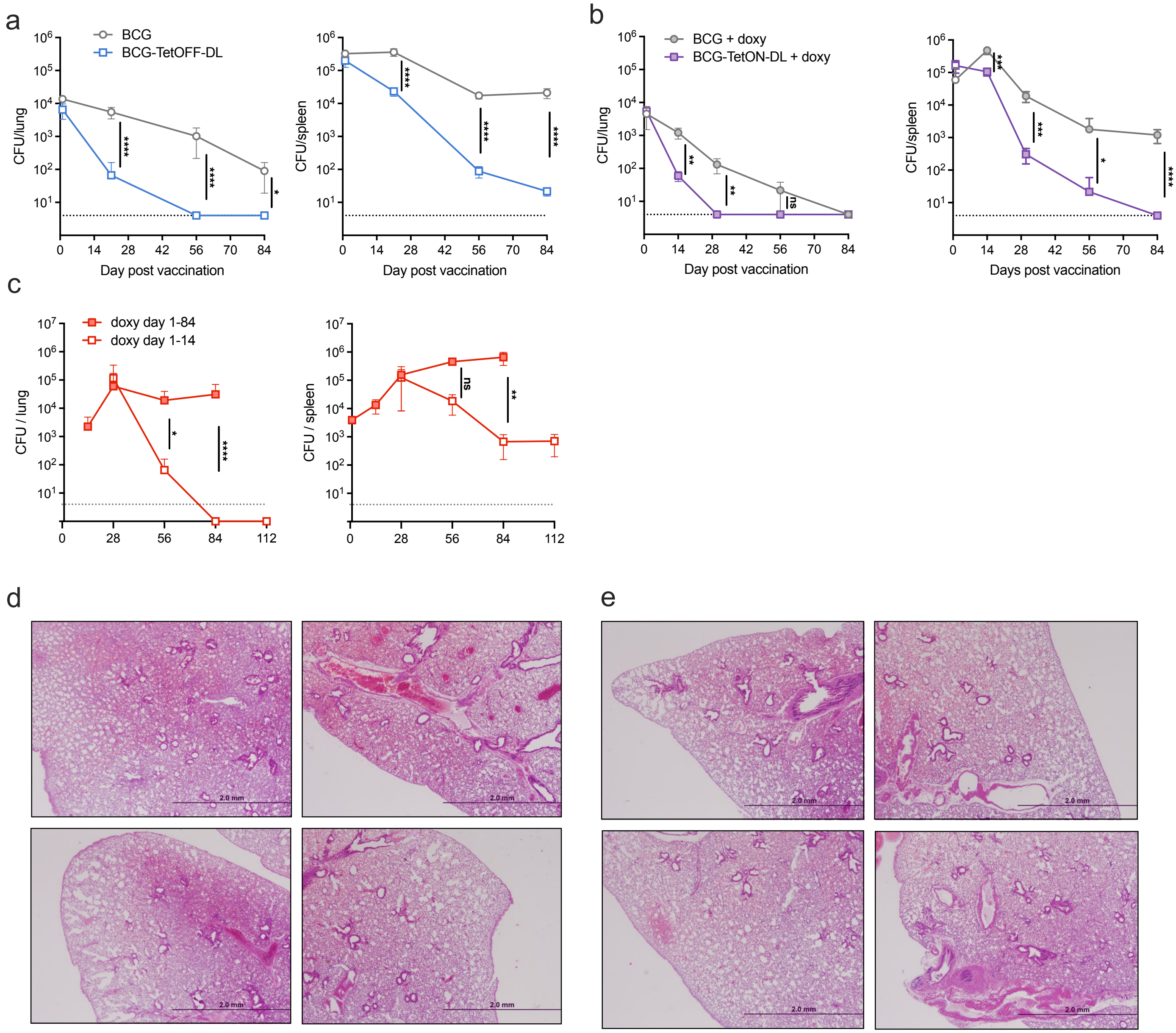

a
BCG
BCG-TetOFF-DL
CFU/lung
CFU/spleen
Day post vaccination
b
BCG + doxy
BCG-TetON-DL + doxy
Days post vaccination
c
doxy day 1-84
doxy day 1-14
CFU / lung
CFU / spleen
d
e
2.0 mm


**Figure S3**

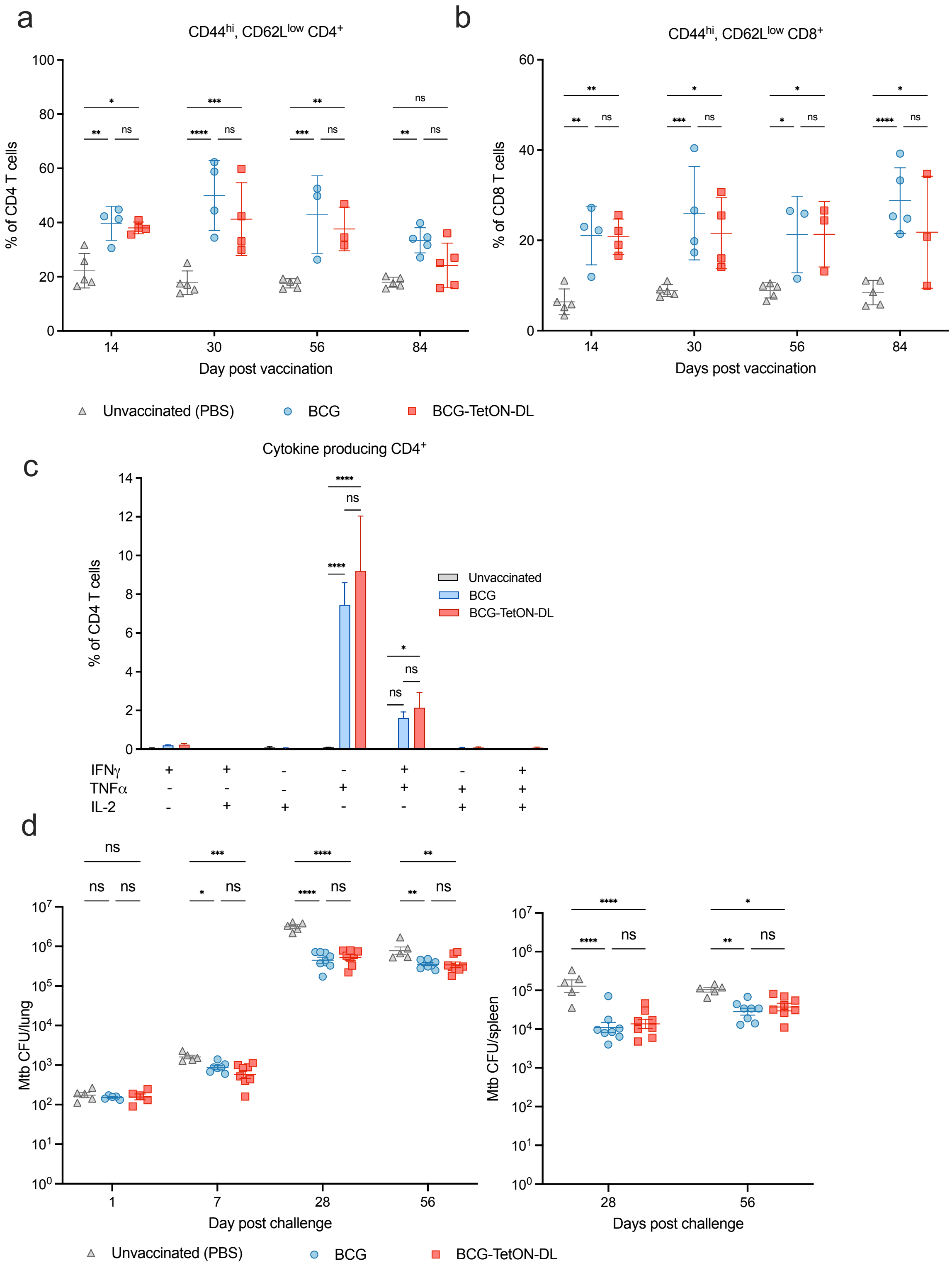
a
CD44hi, CD62Llow CD4+
% of CD4 T cells
Day post vaccination
b
CD44hi, CD62Llow CD8+
% of CD8 T cells
Days post vaccination
Unvaccinated (PBS)
BCG
BCG-TetON-DL
c
Cytokine producing CD4+
% of CD4 T cells
Unvaccinated
BCG
BCG-TetON-DL
IFNγ
TNFα
IL-2
d
Mtb CFU/lung
Day post challenge
Mtb CFU/spleen
Days post challenge
Unvaccinated (PBS)
BCG
BCG-TetON-DL

**Figure S4**

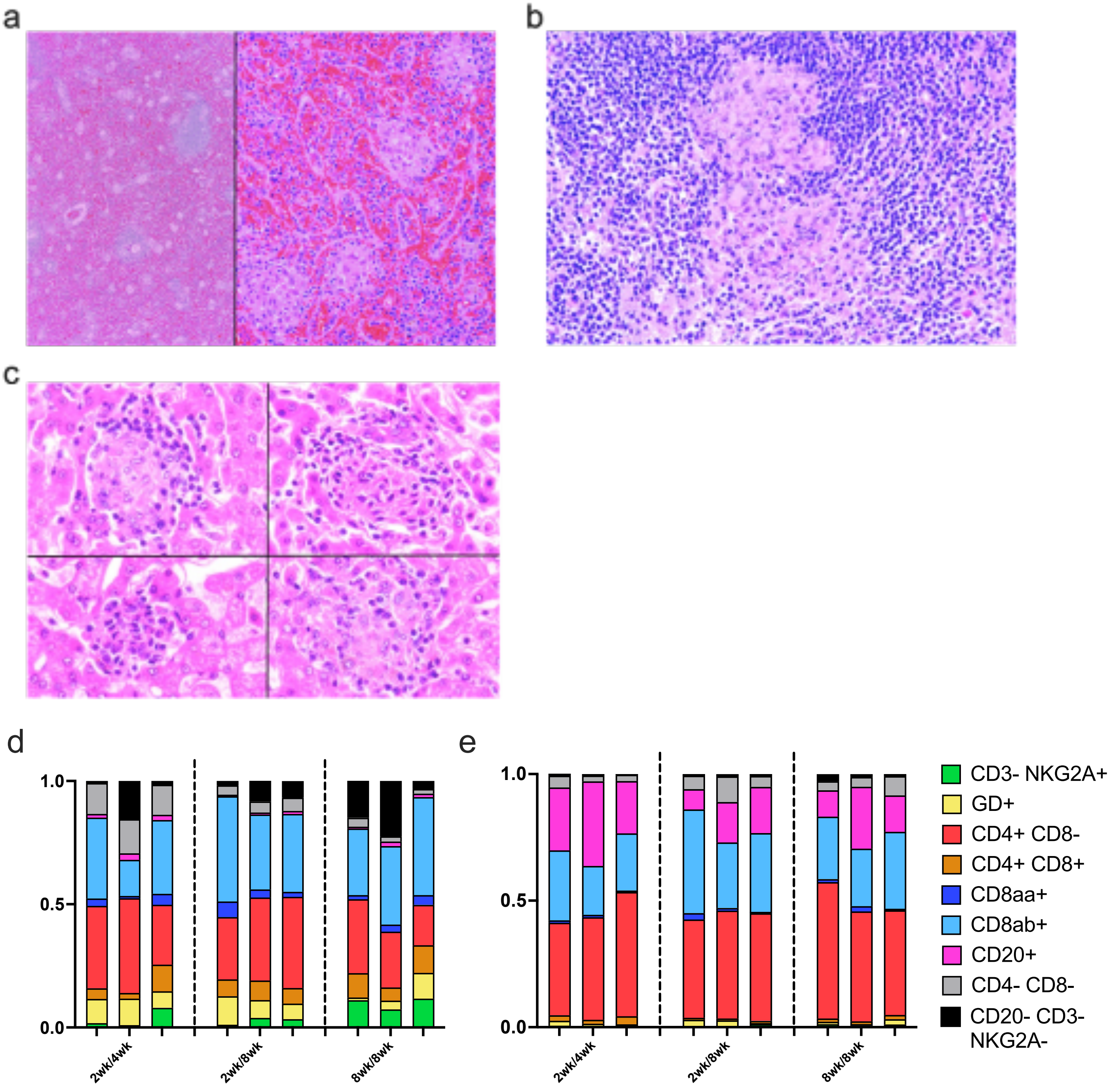
a
b
c
d
e
1.0
0.5
0.0
2wk/4wk
2wk/8wk
8wk/8wk
CD3- NKG2A+
GD+
CD4+ CD8-
CD4+ CD8+
CD8aa+
CD8ab+
CD20+
CD4- CD8-
CD20- CD3- NKG2A-

**Figure S5**

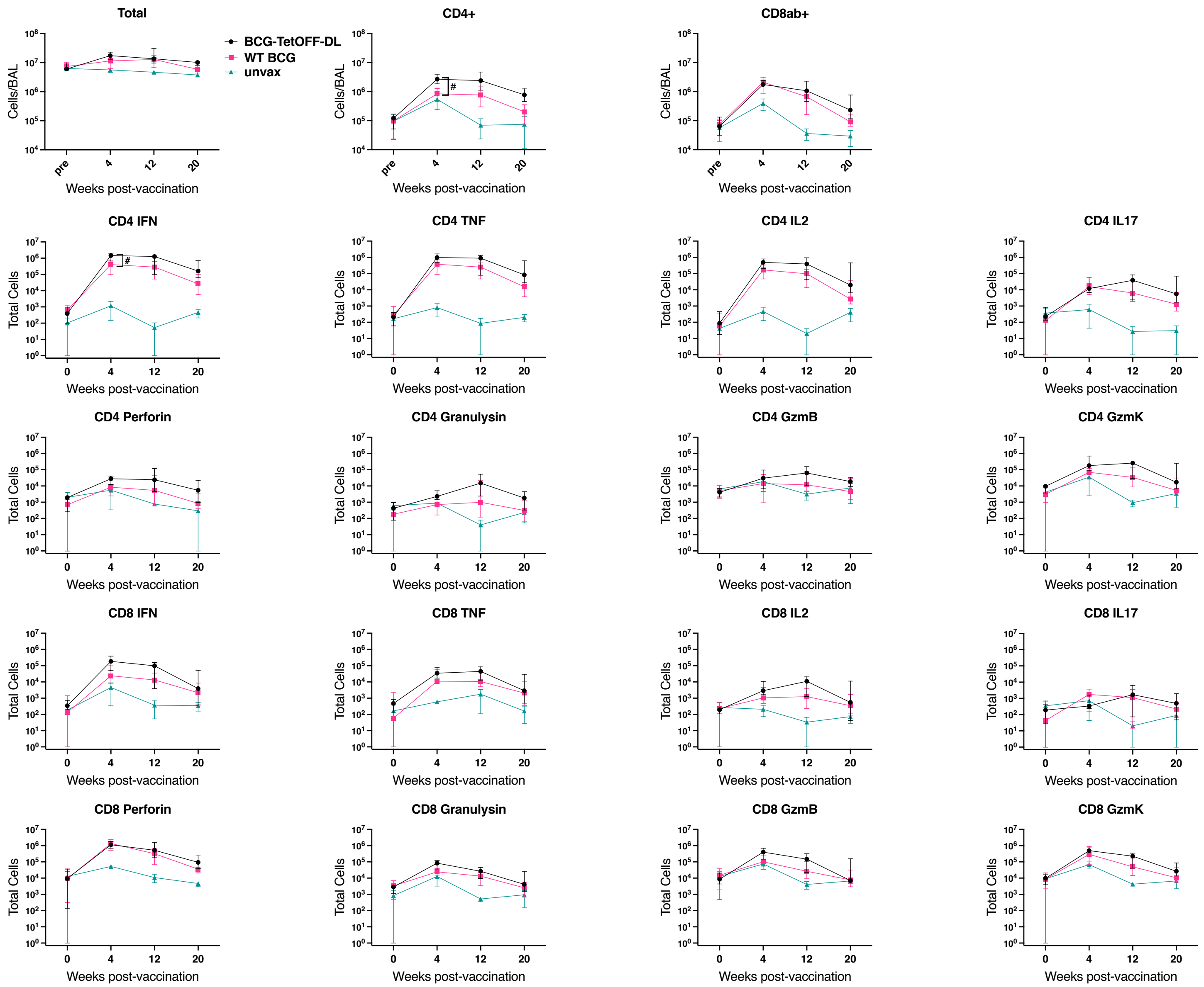
Total
CD4+
CD8ab+
BCG-TetOFF-DL
WT BCG
unvax
Cells/BAL
Weeks post-vaccination
#
CD4 IFN
CD4 TNF
CD4 IL2
CD4 IL17
CD4 Perforin
CD4 Granulysin
CD4 GzmB
CD4 GzmK
CD8 IFN
CD8 TNF
CD8 IL2
CD8 IL17
CD8 Perforin
CD8 Granulysin
CD8 GzmB
CD8 GzmK
Total Cells
pre
0
4
12
20

### Figure S6

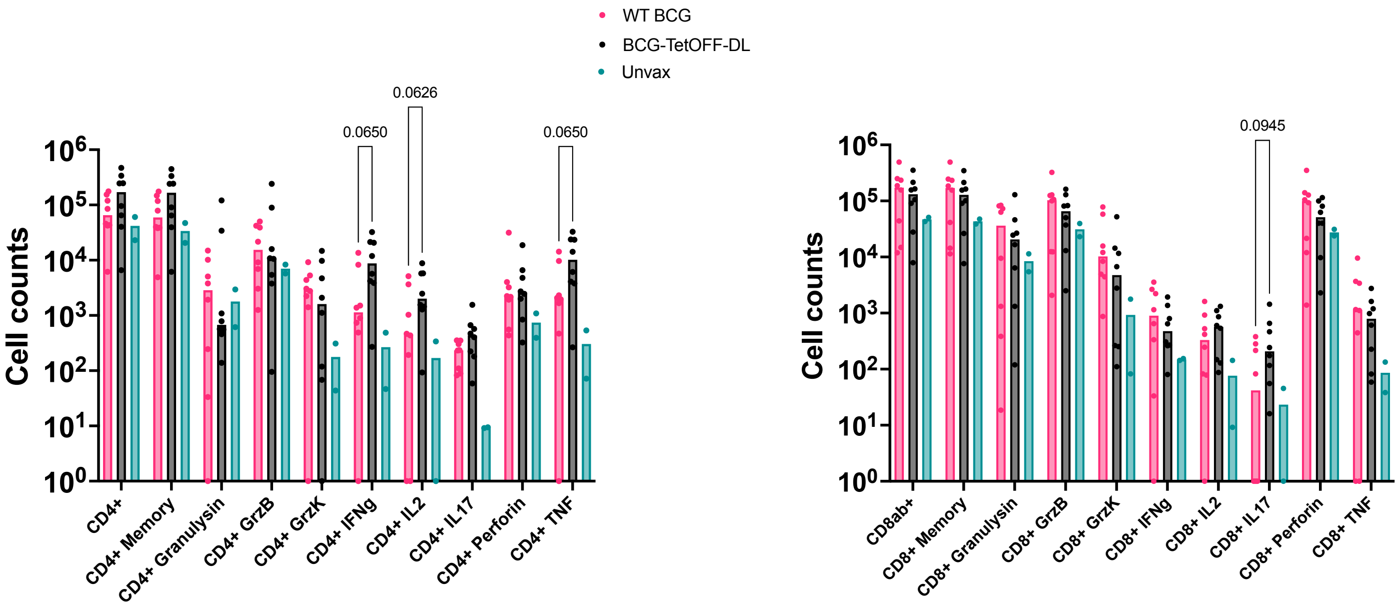

WT BCG
BCG-TetOFF-DL
Unvax
Cell counts
0.0650
0.0626
0.0650
0.0945
CD4+
CD4+ Memory
CD4+ Granulysin
CD4+ GrzB
CD4+ GrzK
CD4+ IFNg
CD4+ IL2
CD4+ IL17
CD4+ Perforin
CD4+ TNF
CD8ab+
CD8+ Memory
CD8+ Granulysin
CD8+ GrzB
CD8+ GrzK
CD8+ IFNg
CD8+ IL2
CD8+ IL17
CD8+ Perforin
CD8+ TNF


**Figure S7**

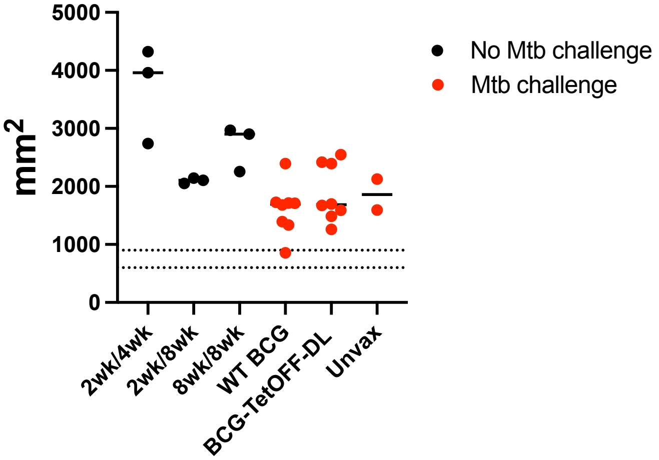
5000
4000
3000
2000
1000
0
mm2
No Mtb challenge
Mtb challenge
2wk/4wk
2wk/8wk
8wk/8wk
WT BCG
BCG-TetOFF-DL
Unvax

**Figure S8**

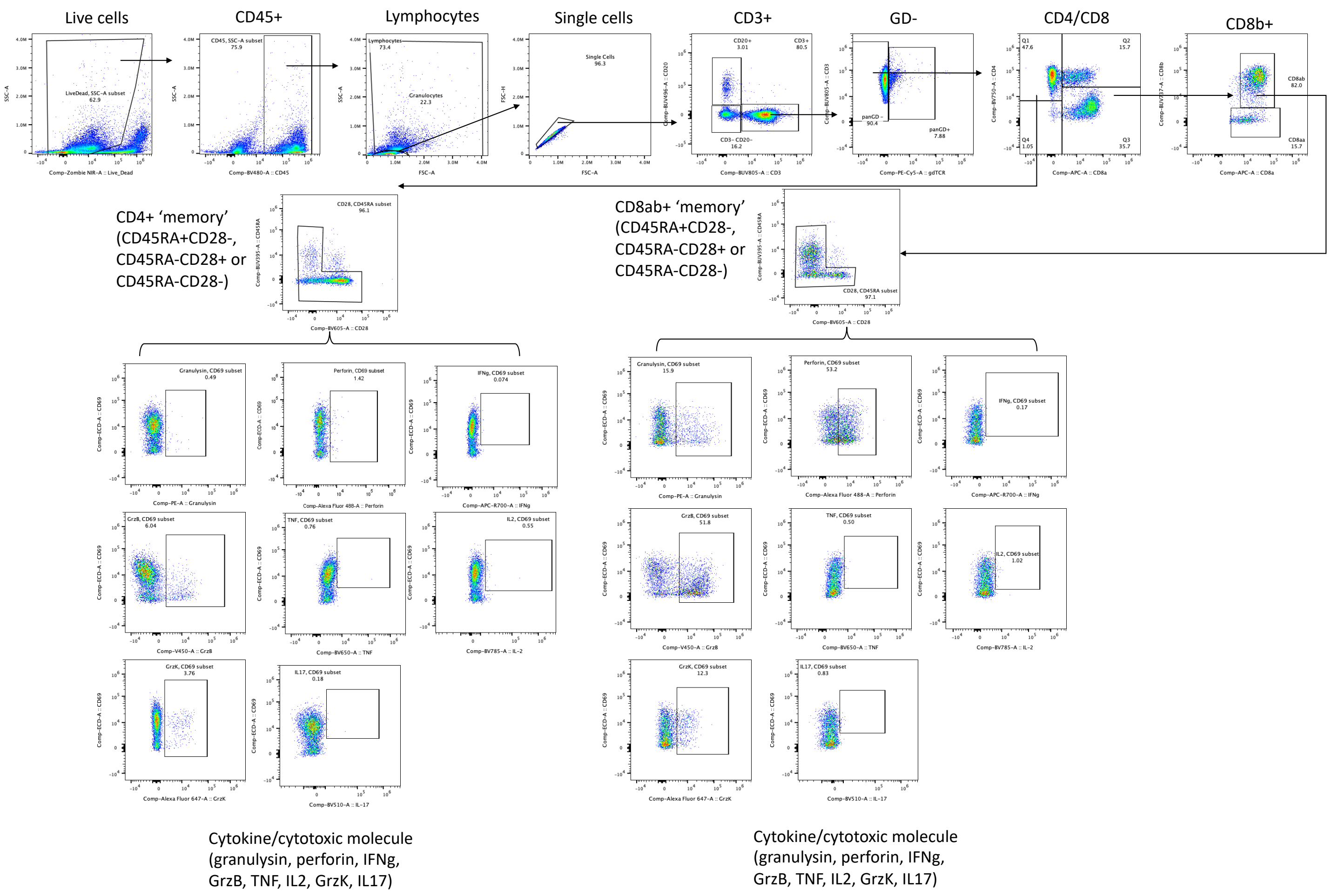
Live cells
CD45+
Lymphocytes
Single cells
CD3+
GD-
CD4/CD8
CD8b+
CD4+ 'memory'
(CD45RA+CD28-,
CD45RA-CD28+ or
CD45RA-CD28-)
CD8ab+ 'memory'
(CD45RA+CD28-,
CD45RA-CD28+ or
CD45RA-CD28-)
Cytokine/cytotoxic molecule
(granulysin, perforin, IFNg,
GrzB, TNF, IL2, GrzK, IL17)
Cytokine/cytotoxic molecule
(granulysin, perforin, IFNg,
GrzB, TNF, IL2, GrzK, IL17)

**Figure S9**

| | Target | Clone | Fluor | Product # | Vendor |
|---|---|---|---|---|---|
| Surface | CD3 | SP34-2 | APC-Cy7 | 624072 | BD Biosciences |
| | CD4 | L200 | BV750 | 747202 | BD Biosciences |
| | CD8a | DK25 | FITC | FCMAB176F | EMD Millipore |
| | CD8b | 2ST8.5H7 | BUV737 | 748324 | BD Biosciences |
| | Pan gd | 5A6.E9 | PE | MHGD04 | Thermo Fischer Scientific |
| | CD16 | 3G8 | BV570 | 302036 | Biolegend |
| | CD159a | Z199 | PE-Cy7 | B10246 | Beckman Coulter |
| | CD45 | D058-1283 | BV480 | 566145 | BD Biosciences |
| | CD28 | CD28.2 | BV605 | 302968 | BD Biosciences |
| | CD45RA | 5H9 | BUV395 | 740315 | BD Biosciences |
| | CD20 | 2H7 | PE-CY5 | 302308 | Biolegend |
| | CD11b | ICRF44 | BUV563 | 741357 | BD Biosciences |
| | CD11c | 3.9 | BV421 | 301628 | Biolegend |
| ICS | CD69 | TP1.55.3 | ECD | 6607110 | Beckman Coulter |
| | Granzyme B | GB11 | V450 | 561151 | BD Biosciences |
| | Granzyme K | G3H69 | AF647 | 566655 | BD Biosciences |
| | IL-17 | BL168 | BV510 | 512330 | Biolegend |
| | TNF | MAb11 | BV650 | 502938 | Biolegend |
| | IFNg | B27 | APC | 506510 | Biolegend |
| | IL-2 | MQ1-17H12 | BV785 | 500348 | Biolegend |
| L/D | Zombie | | NIR | 423106 | Biolegend |

**Table S1**

| NHP | Vendor | Origin | Gender | DOB | BCG Strain | BCG Vax Date | BCG Dose | Doxy Start | Doxy End | Mtb Infectio | Mtb Dose | Necropsy Date | Weeks PI | Grans4wks | PET4wks | Grans8wks | PET8wks | PET12Weeks | NecropsyScore | GransNX | TotalCFU | LungCFU | LNCFU | BodyCFU |
|---|---|---|---|---|---|---|---|---|---|---|---|---|---|---|---|---|---|---|---|---|---|---|---|---|
| 10022 | Bioculture-Mauritius | Mauritius | Male | 7/26/18 | BCG-TetOFFDL | 8/2/22 | 5x10^7 | 8/1/22 | 8/15/22 | 1/4/23 | 9 | 3/27/23 | 11.71429 | 0 | 0 | 0 | 0 | 0 | 8 | 0 | 0 | 0 | 0 | |
| 10122 | Bioculture-Mauritius | Mauritius | Male | 4/25/18 | BCG-TetOFFDL | 8/2/22 | 5x10^7 | 8/1/22 | 8/15/22 | 1/4/23 | 9 | 4/5/23 | 13 | 0 | 0 | 0 | 0 | 0 | 7 | 0 | 0 | 0 | 0 | |
| 10622 | Bioculture-Mauritius | Mauritius | Male | 5/23/14 | WT | 8/2/22 | 5x10^7 | 8/1/22 | 8/15/22 | 1/4/23 | 9 | 4/5/23 | 13 | 5 | 178.119 | 8 | 1214.96702 | 1660.237383 | 17 | 13 | 8550 | 6550 | 2000 | |
| 10722 | Bioculture-Mauritius | Mauritius | Male | 8/6/12 | WT | 8/2/22 | 5x10^7 | 8/1/22 | 8/15/22 | 1/4/23 | 9 | 3/29/23 | 12 | 0 | 0 | 1 | 13.4643538 | 0 | 8 | 0 | 0 | 0 | 0 | |
| 10822 | Bioculture-Mauritius | Mauritius | Male | 10/1/16 | BCG-TetOFFDL | 8/2/22 | 5x10^7 | 8/1/22 | 8/15/22 | 1/4/23 | 9 | 4/3/23 | 12.71429 | 1 | 0 | 1 | 0 | 0 | 9 | 0 | 0 | 0 | 0 | |
| 10922 | Bioculture-Mauritius | Mauritius | Male | 3/1/16 | BCG-TetOFFDL | 8/2/22 | 5x10^7 | 8/1/22 | 8/15/22 | 1/4/23 | 9 | 3/29/23 | 12 | 38 | 12.72893 | 38 | 0 | 0 | 23 | 14 | 0 | 0 | 0 | |
| 11022 | Bioculture-Mauritius | Mauritius | Male | 2/12/18 | WT | 8/2/22 | 5x10^7 | 8/1/22 | 8/15/22 | 1/4/23 | 9 | 4/3/23 | 12.71429 | 5 | 0 | 6 | 6717.6468 | 65104.86686 | 19 | tntc | 2048450 | 2047850 | 600 | |
| 11122 | Bioculture-Mauritius | Mauritius | Male | 1/6/18 | WT | 8/2/22 | 5x10^7 | 8/1/22 | 8/15/22 | 1/4/23 | 9 | 4/10/23 | 13.71429 | 1 | 0 | 1 | 0 | 0 | 9 | 3 | 10038 | 10038 | 0 | |
| 11222 | Bioculture-Mauritius | Mauritius | Male | 4/26/18 | BCG-TetOFFDL | 9/1/22 | 4.9x10^7 | 8/31/22 | 9/13/22 | 1/31/23 | 16 | 4/24/23 | 11.85714 | 1 | 0 | 1 | 0 | 0 | 8 | 0 | 0 | 0 | 0 | |
| 11322 | Bioculture-Mauritius | Mauritius | Male | 8/25/18 | BCG-TetOFFDL | 9/1/22 | 4.9x10^7 | 8/31/22 | 9/13/22 | 1/31/23 | 16 | 4/26/23 | 12.14286 | 0 | 15.56539 | 0 | 0 | 0 | 11 | 0 | 1075 | 0 | 1075 | |
| 11422 | Bioculture-Mauritius | Mauritius | Male | 4/3/18 | WT | 9/1/22 | 4.48x10^7 | 8/31/22 | 9/13/22 | 1/31/23 | 16 | 4/26/23 | 12.14286 | 6 | 0 | 6 | 0 | 0 | 13 | 10 | 5800 | 5400 | 400 | |
| 11522 | Bioculture-Mauritius | Mauritius | Male | 7/1/18 | WT | 9/1/22 | 4.48x10^7 | 8/31/22 | 9/13/22 | 1/31/23 | 16 | 5/10/23 | 14.14286 | 3 | 0 | 3 | 11.8317564 | 0 | 11 | 3 | 705 | 685 | 20 | |
| 11822 | Bioculture-Mauritius | Mauritius | Male | 11/23/16 | BCG-TetOFFDL | 9/1/22 | 4.9x10^7 | 8/31/22 | 9/13/22 | 1/31/23 | 16 | 5/2/23 | 13 | 2 | 0 | 3 | 0 | 0 | 9 | 1 | 30 | 30 | 0 | |
| 11922 | Bioculture-Mauritius | Mauritius | Male | 12/22/16 | BCG-TetOFFDL | 9/1/22 | 4.9x10^7 | 8/31/22 | 9/13/22 | 1/31/23 | 16 | 5/2/23 | 13 | 0 | 0 | 0 | 0 | 0 | 8 | 0 | 0 | 0 | 0 | |
| 12022 | Bioculture-Mauritius | Mauritius | Male | 9/2/18 | WT | 9/1/22 | 4.48x10^7 | 8/31/22 | 9/13/22 | 1/31/23 | 16 | 5/8/23 | 13.85714 | 0 | 0 | 0 | 0 | 0 | 12 | 1 | 0 | 0 | 0 | |
| 12122 | Bioculture-Mauritius | Mauritius | Male | 9/26/18 | WT | 9/1/22 | 4.48x10^7 | 8/31/22 | 9/13/22 | 1/31/23 | 16 | 5/8/23 | 13.85714 | 4 | 0 | 4 | 0 | 0 | 11 | 5 | 270 | 250 | 20 | |
| 12722 | Bioculture-Mauritius | Mauritius | Male | 2/23/16 | none | | | | | 1/4/23 | 9 | 3/27/23 | 11.71429 | 40 | 39853.77 | tntc | 287211.79 | 321798.3491 | 88 | tntc | 1503960 | 1332110 | 136000 | |
| 12822 | Bioculture-Mauritius | Mauritius | Male | 3/14/14 | none | | | | | 1/31/23 | 16 | 4/24/23 | 11.85714 | 6 | 835.3731 | 10 | 24.7 | 0 | 22 | 12 | 20293 | 12533 | 7760 | |
| 15719 | Bioculture-Mauritius | Mauritius | Male | 5/25/14 | none historical | | | | | 12/17/19 | 21 | 03/09/20 | 11.85714 | 19 | 12247.09 | tntc | 110612.275 | 119644.7832 | 75 | tntc | 1359498.29 | 1196783.29 | 162715 | |
| 15819 | Bioculture-Mauritius | Mauritius | Male | 11/25/13 | none historical | | | | | 12/17/19 | 21 | 03/11/20 | 12.14286 | 8 | 5393.503 | 23 | 94418.4916 | 203119.0381 | 56 | tntc | 552445.159 | 502985.159 | 49460 | |
| 15919 | Bioculture-Mauritius | Mauritius | Male | 1/25/12 | none historical | | | | | 12/17/19 | 21 | 03/17/20 | 13 | 12 | 1312.402 | 12 | 776.293776 | 888.2410389 | 23 | 18 | 17740 | 11950 | 5790 | |
| 16019 | Bioculture-Mauritius | Mauritius | Male | 11/25/13 | none historical | | | | | 12/17/19 | 21 | 03/19/20 | 13.28571 | 19 | 9132.703 | 19 | 14718.1518 | 10045.66079 | 37 | 32 | 27504.9491 | 16574.9491 | 10930 | |
| 16119 | Bioculture-Mauritius | Mauritius | Male | 1/25/11 | none historical | | | | | 01/14/20 | 6 | 04/14/20 | 13 | 6 | 978.5377 | tntc | 75438.3824 | 170898.7206 | 70 | tntc | 672398.189 | 637783.189 | 34615 | |
| 16419 | Bioculture-Mauritius | Mauritius | Male | 1/25/13 | none historical | | | | | 01/14/20 | 6 | 04/16/20 | 13.28571 | 8 | 10040.75 | tntc | 68661.0003 | 217106.4084 | 76 | tntc | 199383.539 | 31101.0387 | 168282.5 | |
| 16519 | Bioculture-Mauritius | Mauritius | Male | 1/25/12 | none historical | | | | | 02/14/20 | 4 | 05/12/20 | 12.57143 | 6 | 486.4546 | | | 62842.36694 | 64 | tntc | 7743349.94 | 7648649.94 | 93875 | |
| 16719 | Bioculture-Mauritius | Mauritius | Male | 10/25/14 | none historical | | | | | 02/14/20 | 4 | 05/14/20 | 12.85714 | 1 | 127.9454 | 3 | 536.135708 | 522.973755 | 27 | 13 | 11330 | 2580 | 8750 | |
| 23621 | Bioculture-Mauritius | Mauritius | Male | 12/29/13 | BCG-TetOFFDL | 1/12/22 | 3.74x10^7 | 1/11/22 | 1/26/22 | | | 2/9/22 | | | | | | | 11 | | 160 | 0 | 120 | 160 |
| 23721 | Bioculture-Mauritius | Mauritius | Male | 12/11/15 | BCG-TetOFFDL | 1/12/22 | 3.74x10^7 | 1/11/22 | 1/26/22 | | | 2/9/22 | | | | | | | 7 | | 945 | 615 | 330 | 1041 |
| 23821 | Bioculture-Mauritius | Mauritius | Male | 11/16/12 | BCG-TetOFFDL | 1/12/22 | 3.74x10^7 | 1/11/22 | 1/26/22 | | | 2/7/22 | | | | | | | 23 | | 235 | 175 | 60 | 235 |
| 23921 | Bioculture-Mauritius | Mauritius | Male | 6/19/14 | BCG-TetOFFDL | 1/12/22 | 3.74x10^7 | 1/11/22 | 1/26/22 | | | 3/9/22 | | | | | | | 7 | | 0 | 0 | 0 | 50 |
| 24021 | Bioculture-Mauritius | Mauritius | Male | 6/18/14 | BCG-TetOFFDL | 1/12/22 | 3.74x10^7 | 1/11/22 | 1/26/22 | | | 3/9/22 | | | | | | | 9 | | 0 | 0 | 0 | 0 |
| 24121 | Bioculture-Mauritius | Mauritius | Male | 10/20/14 | BCG-TetOFFDL | 1/12/22 | 3.74x10^7 | 1/11/22 | 3/13/22 | | | 3/14/22 | | | | | | | 6 | | 195 | 145 | 0 | 195 |
| 24221 | Bioculture-Mauritius | Mauritius | Male | 2/5/15 | BCG-TetOFFDL | 1/12/22 | 3.74x10^7 | 1/11/22 | 3/6/22 | | | 3/7/22 | | | | | | | 7 | | 25 | 0 | 0 | 25 |
| 24321 | Bioculture-Mauritius | Mauritius | Male | 6/26/14 | BCG-TetOFFDL | 1/12/22 | 3.74x10^7 | 1/11/22 | 3/8/22 | | | 3/9/22 | | | | | | | 9 | | 85 | 85 | 0 | 85 |
| 24421 | Bioculture-Mauritius | Mauritius | Male | 6/8/13 | BCG-TetOFFDL | 1/12/22 | 3.74x10^7 | 1/11/22 | 1/26/22 | | | 3/14/22 | | | | | | | 24 | | 15 | 15 | 0 | 15 |

| Variable | Description |
|---|---|
| **NHP** | Non-Human Primate ID |
| **Vendor** | Animal Vendor |
| **Origin** | Animal Origin |
| **Gender** | Gender |
| **DOB** | Date of Birth |
| **BCG Strain** | BCG Strain: WT= wild type BCG Pasteur, BCG-TetOFFDL = BCG Dual lysin Pasteur kill switch strain, none = Unvaccinated, none historical = Unvaccinated control from previous study |
| **BCG Vax Date** | Date of BCG Vaccination |
| **BCG Dose** | Estimated BCG Dose |
| **Doxy Start** | Doxycycline Start Date |
| **Doxy End** | Doxycycline End Date |
| **Mtb Infection** | Date of Mtb Infection |
| **Mtb Dose** | Estimated Mtb Dose |
| **Necropsy Date** | Date of Necropsy |
| **Weeks PI** | Weeks Post Mtb Infection |
| **Grans4wks** | Number of Granulomas at 4 Weeks Post-Infection |
| **PET4wks** | Total Lung FDG Activity at 4 Weeks Post-Infection |
| **Grans8wks** | Number of Granulomas at 8 Weeks Post-Infection (tntc = too numerous to count or TB Pneumonia) |
| **PET8wks** | Total Lung FDG Activity at 8 Weeks Post-Infection |
| **PET12Weeks** | Total Lung FDG Activity at 12 Weeks Post-Infection |
| **NecropsyScore** | Gross Pathology Score |
| **GransNX** | Number of Granulomas Found at Necropsy (tntc = too numerous to count or TB Pneumonia) |
| **TotalCFU** | Total CFU |
| **LungCFU** | Lung CFU |
| **LNCFU** | Thoracic Lymph Node CFU |
| **BodyCFU** | Estimated Total Body CFU |

**Table S2**
